# Neural xenografts contribute to long-term recovery in stroke via molecular graft-host crosstalk

**DOI:** 10.1101/2024.04.03.588020

**Authors:** Rebecca Z. Weber, Beatriz Achón Buil, Nora H. Rentsch, Patrick Perron, Stefanie Halliday, Allison Bosworth, Mingzi Zhang, Kassandra Kisler, Chantal Bodenmann, Kathrin J. Zürcher, Daniela Uhr, Debora Meier, Siri L. Peter, Melanie Generali, Shuo Lin, Markus A. Rüegg, Roger M. Nitsch, Christian Tackenberg, Ruslan Rust

## Abstract

Stroke is a leading cause of disability and death due to the brain’s limited ability to regenerate damaged neural circuits. To date, stroke patients have only few therapeutic options and are often left with considerable disabilities. Induced pluripotent stem cell (iPSC)-based therapies are emerging as a promising therapeutic approach for stroke recovery. In this study, we demonstrate that local transplantation of iPSC-derived neural progenitor cells (NPCs) improves long-term brain tissue repair responses and reduces neurological deficits after cerebral ischemia in mice. Using *in vivo* bioluminescence imaging and *post-mortem* histology, we show long-term graft survival over the course of five weeks and preferential graft differentiation into mature neurons without signs of pluripotent residuals. Transplantation of NPCs led to a set of brain tissue repair responses including increased vascular sprouting and repair, improved blood-brain barrier integrity, reduced microglial activation, and increased neurogenesis compared to littermate control animals receiving sham transplantation. Employing deep learning-assisted behavior analysis, we found that NPC-treated mice displayed improved gait performance and complete fine-motor recovery in the horizontal ladder rung walk, five weeks post-injury. To dissect the molecular graft composition and identify graft-host interactions, single nucleus profiling of the cell transplants and host stroke tissue was performed. We identified graft differentiation preferentially towards neurons with GABAergic and glutamatergic phenotypes in similar proportions, with the remaining cells acquiring astrocyte and NPC-like phenotypes. Interaction between graft and host transcriptome indicated that GABAergic cell grafts were primarily involved in graft-host communication through the regeneration-associated neurexin (NRXN), neuregulin (NRG), neural cell adhesion molecule (NCAM) and SLIT signalling pathways. In conclusion, our study reveals that transplanted iPSC-derived NPCs primarily differentiate into GABAergic neurons contributing to long-term recovery and further delineates the regenerative interactions between the graft and the stroke-injured host tissue.

## Introduction

Ischemic stroke is a major cause of disability and death worldwide, affecting 1 in 4 adults in their lifetime^1,2^. Current treatment options including intravenous thrombolysis and mechanical thrombectomy are constrained by a narrow therapeutic time window and potential complications, leaving 50% of patients with remaining long-term disabilities^3^. Stem cell therapies have shown beneficial effects in stroke recovery in various animal models by immune modulation^4,5^, neurogenesis^6,7^, angiogenesis^5,6^, blood-brain barrier repair^8^ and neural circuit restoration^9–12^ in the stroke-injured brain. While multiple cell sources such as mesenchymal stem cells (MSCs)^13–15^, embryonic stem cells (ESCs)^16^, or fetal-derived neural stem/progenitor cells (NSCs/NPCs)^5,17^ have shown encouraging preclinical results, their efficacy could not be confirmed in recent large clinical trials with stroke patients^18,19^.

Recent advances in genetic engineering and cell culture protocols increased the therapeutic potential of induced pluripotent stem cell (iPSC)-derived NPCs, which have unique features such as scalability, reduced ethical concerns, neural differentiation potential, and patient-specific adaptability^20,21^. For instance, xeno- and transgene-free protocols have been developed for hiPSC-NPCs in accordance with good manufacturing practices (GMP)^22^. To reduce the risk of tumor formation, the safety profile of iPSC-derived cell sources has been recently improved using genetic safety switch systems^22,23^. Previous studies showed that transplantation of iPSC-derived NPCs can release neurotrophic factors, form new connections within the damaged neural circuits^9,24^ and repair functional deficits in stroke animals^25,26,10,8,7,27^. However, these studies only focused on selected aspects of regeneration, performed a histology-based graft characterisation, and did not analyse the molecular crosstalk between graft and host. The molecular graft-host interaction is largely unknown^28^ and to our knowledge has not been assessed at single-cell resolution.

Here we demonstrate the long-term survival and integration of NPCs into stroke-injured mouse brains. Mice that received NPC grafts, locally transplanted in the injured brain tissue adjacent to the infarct region, exhibited regeneration-associated tissue responses five weeks post-stroke including reduced inflammation, enhanced angiogenesis, and increased neuro- and axonogenesis in NPC-treated mice, compared to their sham-treated littermates. The observed anatomical improvements were associated with increased functional recovery determined by deep learning-assisted behavioral assessments. Histological and molecular characterization of the cell graft revealed a preferential differentiation from NPCs to GABAergic neurons. Molecular profiling of single cell graft and surrounding host cells suggests a potential graft-host crosstalk, where GABAergic cell grafts may induce tissue regeneration through Neurexin (NRXN), Neuregulin (NRG), Neural Cell Adhesion Molecule (NCAM), and SLIT signaling pathways.

## Results

### Transplantation of hiPSC-NPCs seven days after stroke results in long-term graft survival

To generate a cell graft with translational potential we differentiated human iPSCs into neural progenitor cells (hiPSC-NPCs, hereafter called “NPCs”) under transgene- and xeno-free conditions that can be smoothly adapted to GMP-grade for clinical applications as previously described^22^. The cell source has previously been characterized in-depth to spontaneously differentiate into astrocytes and functional, mature neurons and is compatible with safety switches for safe *in vivo* applications^22^. First, we confirmed the presence of canonical NPC markers including Nestin and Pax6 and the absence of pluripotent markers such as Nanog in NPCs (**Fig. 1A, Suppl. Fig. 1**). Successful spontaneous differentiation upon growth factor withdrawal over a 4-week period was validated by upregulation of differentiation markers such as neuronal βIII-Tubulin and astrocytic S100β (**Fig. 1B**).

**Fig. 1:**
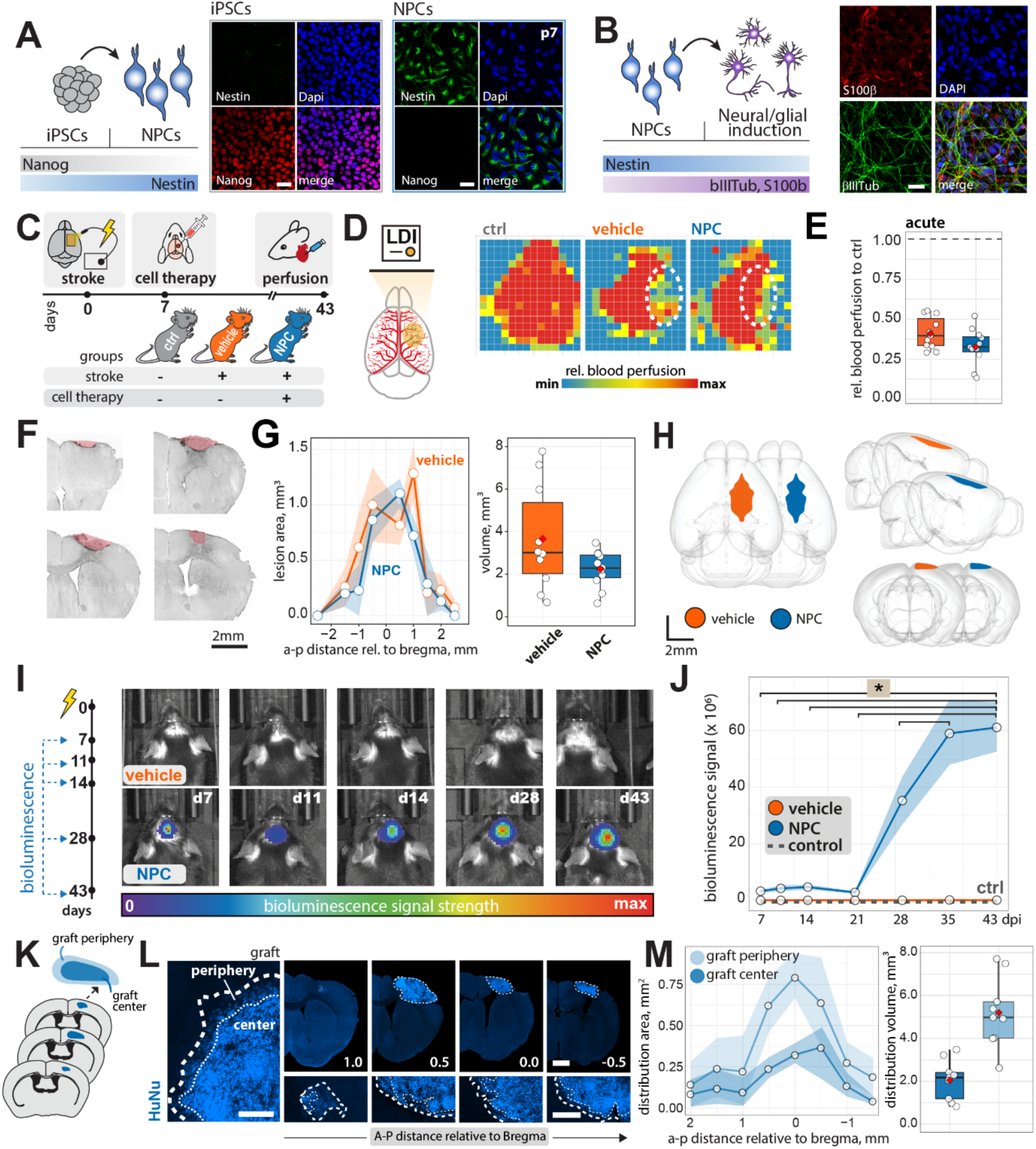
Generation of hiPSC-NPCs, stroke induction and transplantation. (A) Left: Generation of hiPSC-NPCs. Right: iPSCs and NPCs (passage 7) stained for Nanog, iPSC marker, and Nestin, a NPC marker. Scale bar: 50um. (B) Left: Neural differentiation of NPCs. Right: Differentiated NPCs, at d26 after differentiation (upper row), stained for βIII-Tubulin, a marker for mature neurons, S100β, a marker for astrocytes, and DAPI. Scale bar: 50um. (C) Schematic representation of experimental design. (D) Brain perfusion levels acquired through Laser-Doppler imaging (LDI). (E) Relative blood perfusion of the right hemisphere compared to baseline recorded immediately after stroke induction (acute) and before sacrifice (43 dpi). (F) Coronal brain sections used to estimate stroke area. The dotted red line indicates the stroke-lesioned brain area. Scale bar: 2mm (G) Quantification of stroke infarction size. Left: Lesion area plotted against the anterior-posterior (a-p) distance relative to bregma (mm). Right: boxplot displaying lesion volumes (mm^3^) of both treatment groups. (H) Schematic representation of a 3-D mouse brain model depicting stroke infarction sizes. Scale bar: 2mm. (I) Longitudinal analysis of cell survival following NPC transplantation using bioluminescence imaging. (J) Bioluminescence signal intensity is represented as the number of photons per second per cm^3^ of sr x10^6^ over a time course of 35 days. Shown significance levels refer to comparisons between days. (K) Schematic and immunofluorescent representation depicting graft core (dark blue) and graft periphery (light blue). HuNu was used to visualize transplanted cells. Scale bar: 1mm. (L) Brain sections stained for HuNu arranged in anterior-to-posterior (a-p) order. Scale bar: 2mm. (M) Quantification of graft core and graft peripheral area. Left: graft area (mm^2^) plotted against the anterior-posterior (a-p) distance relative to bregma (mm). Right: boxplot displaying mean graft volume (mm^3^) of transplanted animals. Data are shown as mean distributions where the red dot represents the mean. Boxplots indicate the 25% to 75% quartiles of the data. For boxplots: each dot in the plots represents one animal. Line graphs are plotted as mean ± sem. Significance of mean differences was assessed using a paired t-test (baseline vs. stroke) or an unpaired t-test (vehicle vs. NPC). In E-J, n=11 mice per group; in M, n=9 animals per group. Asterisks indicate significance: *p < 0.05.

All experimental animal groups underwent a photothrombotic stroke or sham surgery in the right sensorimotor cortex. At 7 days post-injury (dpi), one group of mice received a local transplantation of NPCs expressing a dual-reporter (red firefly luciferase and eGFP, rFluc-eGFP, **Suppl. Fig. 2**), while the other groups (vehicle and control) were subjected to a sham transplantation (**Fig. 1C**). Cerebral blood perfusion was assessed in all animals by Laser-Doppler Imaging (LDI) to confirm stroke induction after surgery (**Fig. 1D**). LDI data demonstrated a 60-70% reduction in cerebral blood flow in both groups compared to non-stroked animals 30 min after stroke induction (vehicle: −61%, NPC: −69% **Fig. 1E**). There was no significant change in perfusion between treatment and control group (p > 0.05). Five weeks after stroke induction, infarcted tissue extended from +2.5 to –2.5 mm anterior-posterior relative to bregma (**Fig. 1F**). A non-significant decrease in lesion volume was observed in mice that received NPCs compared to the vehicle group (NPC: 2.24 mm^3^; vehicle: 3.66 mm^3^, p=0.09, **Fig. 1G, H**).

Transplanted cell survival was confirmed over a 35-day time course using luciferase bioluminescence imaging (**Fig. 1I**). The bioluminescence signal remained relatively constant during the first two weeks after implantation (dpi 7-21), followed by a significant increase in bioluminescent signal until dpi 35 (p < 0.05; **Fig. 1J**). No increase in bioluminescence signal has been observed between 35 and 43 dpi (p = 0.99). As expected, no signal was detected in vehicle and sham control treatment groups that had not received cells.

To quantify cell graft size and migration, confocal microscopy images of serial brain sections stained with an antibody against human nuclei (HuNu) were evaluated. As NPCs can migrate within the brain^29^, we delineated the graft distribution in two distinct areas: the “graft center”, defined as the region with densely populated cells (>3x HuNu signal to background), and the “graft periphery”, where cell density was reduced (HuNu signal threshold between >1.5x and <3x to background; **Fig. 1K**). Areas of graft center and graft periphery were measured for all brain sections and identified with their stereotaxic coordinates (**Fig. 1L**). The largest distribution of peripheral graft cells was observed at bregma (graft distribution area: center = 0.32 mm^2^; periphery = 0.8 mm^2^); only minor migration of cells was found on the most anterior and posterior brain sections (0.12 mm^2^ and 0.25 mm^2^, respectively) (**Fig. 1M**). Together, this data indicates that NPCs grafts are capable of surviving for extended periods after implantation and are likely to migrate towards neighboring cortical regions.

### NPC transplantation affects microglia response

After stroke, resting microglia react to the inflammatory stroke milieu and change their morphology from highly ramified into a more amoeboid type, depending on the distance to the lesion site (**Fig. 2A**). We therefore quantified three morphological features (branching, circularity, ramification) in addition to Iba1 fluorescence intensity to study the effect of NPC therapy on microglia activation (**Fig. 2B**). First, relative immunoreactivity levels of Iba1 were compared between treatment groups in the stroke core of the ipsilateral hemisphere and a corresponding region of interest (ROI) in the contralesional hemisphere for each mouse. Overall, we found an increased immunoreactivity of Iba1 in the ipsilateral cortex, with a strong signal in the stroke core, compared to the contralesional cortex in both groups. However, mice that received NPC exhibited a significant 32% reduction in Iba1 immunoreactivity within the stroke core relative to the vehicle group (Iba1 fluorescence intensity: NPC: +280%, vehicle: +410%, p = 0.02, **Fig. 2C**). As expected, no significant differences in Iba1 immunoreactivity were detected between the groups on the contralesional cortex (p = 0.42, **Fig. 2C, Suppl. Fig. 3A**). Microglial branching index, a measure of microglial branching complexity, and the ramification index, a measure of the ratio of the cell’s perimeter to its area, was significantly reduced in the ischemic border zone (IBZ) compared to their respective contralateral ROIs. However, mice that received NPC therapy showed increased numbers in both, the branching index (vehicle: 19.1±23.8, NPC: 29±37.7, p<0.05) and the ramification index (vehicle: 1.09±0.16, NPC: 1.37±0.35, p<0.001) compared to vehicle animals (**Fig. 2D**). Importantly, no significant differences in branching index and ramification index were detected between the groups on the contralesional cortex (p>0.05), and there were no significant differences in circularity between the treatment groups in the contralesional and the ipsilesional area of interest (p>0.05). Elevated levels of GFAP immunoreactivity, indicating glial scar formation, were observed in the IBZ compared to the contralesional cortex in both groups (**Fig. 2E, Suppl. Fig. 3A**). The increase in GFAP immunoreactivity was comparable between both treatment groups (GFAP fluorescence intensity: NPC: +1150%, vehicle: +1350%, p = 0.65, **Fig. 2F**).

**Fig. 2:**
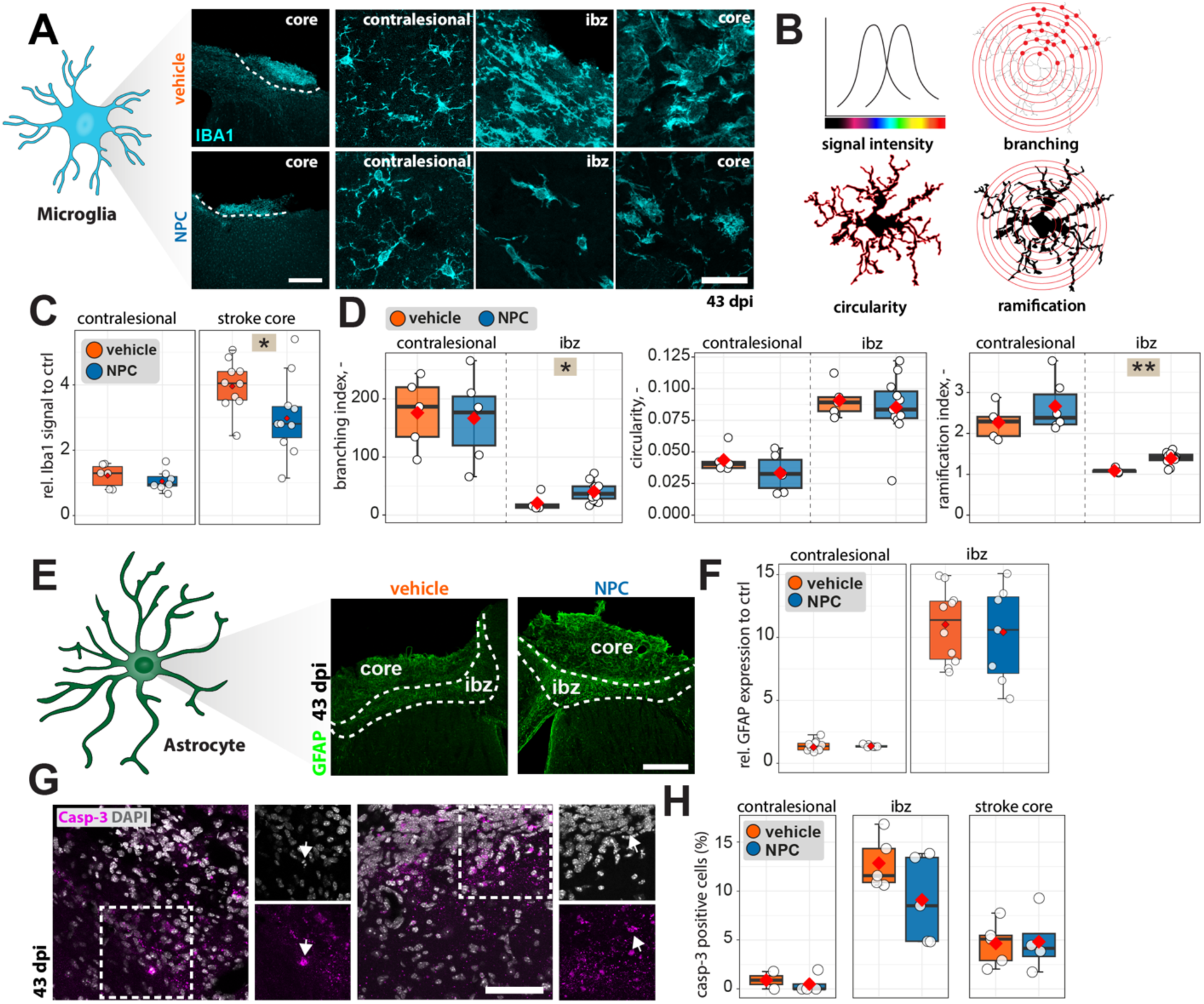
Anatomical changes after stroke induction in NPC- and vehicle receiving mice. (A) Immunofluorescent staining of coronal brain sections with Iba1. Scale bar: 50 µm (left) and 5 µm (right) (B) Scheme depicting the analysed parameters; Iba1 signal intensity and microglia morphology (branching, circularity, ramification). (C) Quantification of Iba1 immunoreactivity on the unaffected contralesional hemisphere and in the stroke core. Data is relative to unstroked control. (D) Quantification of microglia morphology parameters (i) branching index, (ii) circularity and (iii) ramification index. Data is relative to unstroked control. (E) Immunofluorescent staining with GFAP. Scale bar: 50 µm. The ischemic border zone (IBZ) and stroke core are shown to illustrate the stroke-lesioned area. (F) Quantification of GFAP immunoreactivity in the ischemic border zone. (G) Representative histological overview of caspase-3 and Dapi. Scalebar: 10um. (H) Quantification of casp-3 positive cells in the unaffected contralesional hemisphere, the ischemic border and the stroke core zone. Data is relative to all counted Dapi-positive cells. Data are shown as mean distributions where the red dot represents the mean. Boxplots indicate the 25% to 75% quartiles of the data. For boxplots: each dot in the plots represents one animal. Significance of mean differences was assessed using an unpaired t-test (vehicle vs. NPC). In C, n=10 mice per group. In D and H, n=5 animals per group. In F, n=10 (vehicle) and n=7 (NPC) mice per group. Asterisks indicate significance: *p < 0.05, **p < 0.01, ***p < 0.001.

To explore apoptopic cells in the ischemic border zone and the stroke core, capsase-3 immunohistochemical stainings were performed at 43 days after stroke induction (**Fig. 2G**). Numerous caspase-3^+^ cells were found in the ischemic border zone, as well as in the stroke core, compared to the uninjured contralesional side (**Fig. 2H**). However, we did not find any significant differences in caspase-3^+^ counts between NPC and vehicle treatment groups. To further characterize apoptotic cells and investigate potential differences between the two treatment conditions, we performed co-labeling with markers for endothelial cells (CD31, **Suppl. Fig. 3B**) and mature neurons (NeuN, **Suppl. Fig. 3C**). In the IBZ, 18.8 ± 6% of caspase-3^+^ cells co-expressed CD31 in the NPC group, compared to 13.1 ± 4% in the vehicle group (p=0.46). In the stroke core, 24.8 ± 9% of caspase-3^+^ cells were CD31^+^ in the NPC group versus 18.3 ± 6% in the vehicle group (p=0.57). Co-labeling with NeuN revealed that in the IBZ, 27 ± 7% of caspase-3^+^ cells were NeuN^+^ in the NPC group and 17.5 ± 4% in the vehicle group (p=0.26). In the stroke core, we observed 15.1 ± 3% NeuN/caspase-3 double-positive cells in the NPC group and 23.9 ± 8% in the vehicle group, indicating a trend toward a higher proportion of apoptotic neurons in vehicle-treated animals, although this difference did not reach statistical significance (p=0.34).

These results suggest that NPC therapy reduces microglial activation but does not affect glial scarring or apoptosis.

### Increased neurite outgrowth and endogenous neurogenesis in NPC-receiving animals

The immunoreactivity levels of neurofilament light (NF-L) and heavy chain (NF-H), major proteins of the neuronal cytoskeleton, were quantified to determine overall axon regeneration in the stroke core and the surrounding IBZ and were normalized to the unaffected contralesional cortex (**Fig. 3A, B**). We observed significantly higher levels of NF-H in the IBZ of the NPC group (NPC: +48%, p < 0.05) compared to the vehicle group, but no differences in in the stroke core (**Fig. 3C**). For NF-L, we detected a significant increase in relative fluorescence intensities in the stroke core of the NPC group (NPC: +63%, p < 0.005) compared to animals that did not receive cell therapy. No changes have been observed between the groups for NF-L expression in the IBZ (**Fig. 3D**).

**Fig. 3:**
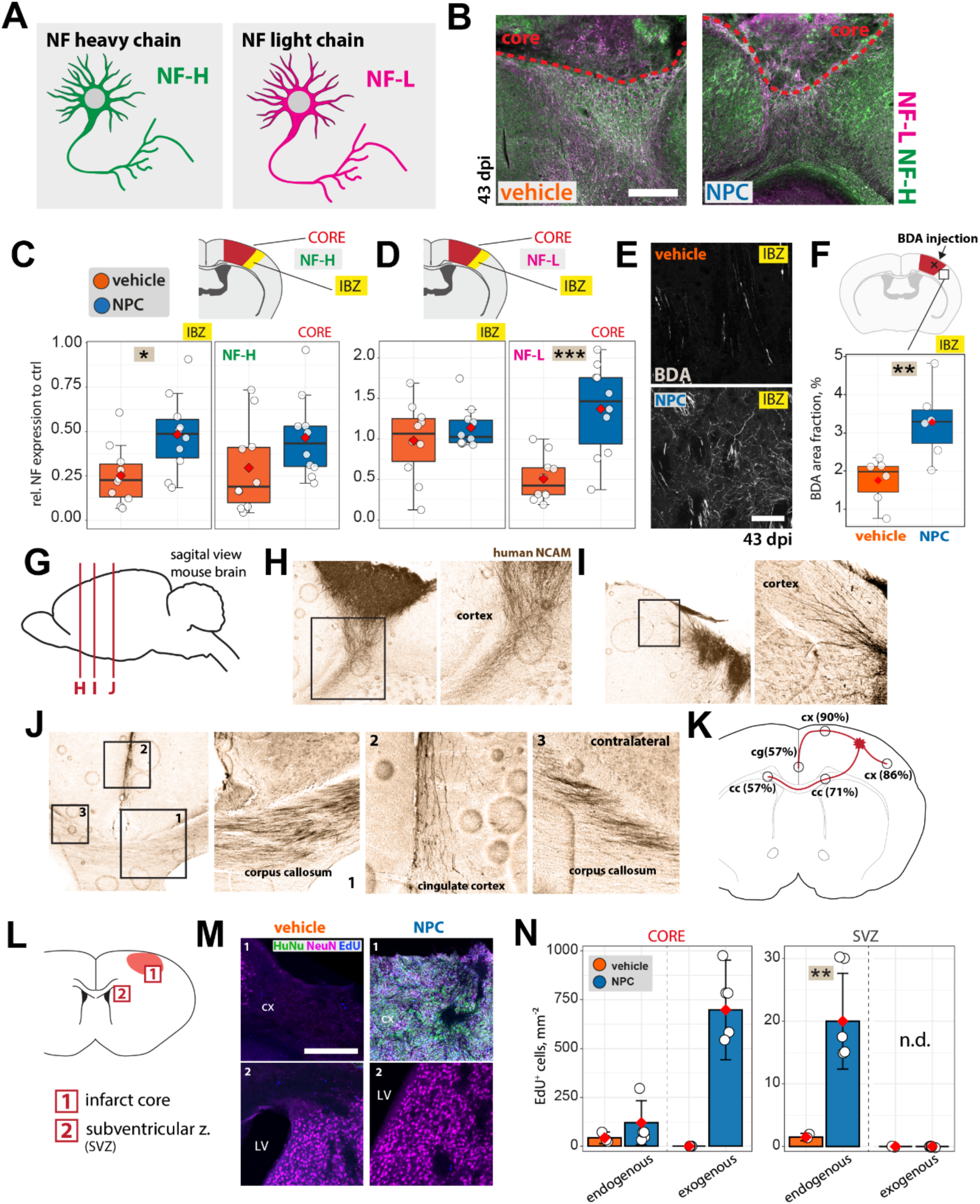
Endo- and exogenous neurite outgrowth and neurogenesis. (A) Representative histological overview of neurofilament heavy chain (NF-H, green) and light chain (NF-L, magenta). (B) Immunofluorescent staining of a coronal brain section with NF-H and NF-L (merge (top) and single channels (bottom). Scale bar: 50 µm (top), 10 µm (bottom). (C) Quantification of relative neurofilament light chain and (D) neurofilament heavy chain signal in the ischemic border and the stroke core zone. Data is normalized to unaffected contralesional site. (E) Representative confocal images of biotinylated dextran amine (BDA) staining in the ipsilateral ischemic border zone of vehicle- and NPC receiving animals. (F) Quantification of BDA-positive fibers relative to BDA-labelled cell bodies. Scale bar = 10µm. (G) Schematic illustration of brain sections corresponding to the following images. (H-J) Representative images of hNCAM staining at different brain levels. (K) The percentage of animals in which transplanted NPCs extended neurites to various brain regions. (L) Illustration showing the two brain areas (infarct core, subventricular zone) analysed for EdU-positive cells. (M) Immunofluorescent staining of a coronal brain section with EdU, NeuN and HuNu and (N) quantification of EdU+ cell count relative to all counted Dapi+ cells. Scale bar = 50µm. Data are shown as mean distributions where the red dot represents the mean. Boxplots indicate the 25% to 75% quartiles of the data. For boxplots: each dot in the plots represents one animal. Significance of mean differences was assessed using an unpaired t-test (vehicle vs. NPC). In C-E, n=10 mice per group. In F, n=6 per group. In N, n=5 mice per group. Asterisks indicate significance: *p < 0.05, **p < 0.01, ***p < 0.001.

To visualize axons, the anterograde axonal tracer biotinylated dextran amine (BDA) was microinjected into the ischemic border zone in ipsilesional cortex adjacent to the stroke area of NPC- and vehicle-treated mice one week prior to sacrifice. Axonal density was quantified using fluorescence intensities normalized to BDA-labelled cell bodies and compared between NPC- and vehicle-receiving animals (**Fig. 3E**). NPC-grafted animals showed increased axonal connection density within the IBZ, compared to the vehicle group (1.83% ± 0.342%; NPC: 3.4% ± 0.36%, p = 0.0055, **Fig. 3F**). Together this data indicates that engrafted NPCs promote neuro- and axonogenesis in and around the infarct region.

Neurite extensions derived from the NPC graft were evaluated using human-specific neural cell adhesion molecule (hNCAM) staining (**Fig. 3G-J**). Among animals receiving cell transplants, 71% exhibited axonal extensions to the ipsilateral corpus callosum (cc), and 57% exhibited their neurites along the cc to the contralateral hemisphere **(Fig. 3K)**. In 57% of the animals that received NPC grafts, we observed axonal extensions to the cingulate cortex. Notably, in 90% of all transplanted animals, cells extended their neurites along the ipsilateral cortex to the primary and secondary motor cortex and the primary somatosensory cortex.

NPC transplantation has previously been shown to promote endogenous neurogenesis in the subventricular zone (SVZ) and subsequent migration to the ischemic lesion site^30^. We therefore assessed if NPC-transplantation positively influenced endogenous neurogenesis in the ischemic cortex. We injected NPC- and vehicle-receiving animals with the nucleoside analog 5-ethynyl-2′-deoxyuridine (EdU) at day 7 following transplantation and, after sacrifice, identified EdU^+^/NeuN^+^ double-positive neurons (endogenous neurogenesis) and EdU^+^/NeuN^+^/HuNu^+^ tripple-positive neurons (exogenous neurogenesis) on a subset of coronal brain sections in the infarct core and the subventricular zone (**Fig. 3L, M**). While we could not detect any differences in endogenous neurogenesis in the ischemic core, we found a significant increase in EdU^+^/NeuN^+^ double-positive neurons in the SVZ in NPC-treated animals compared to the vehicle group (vehicle: 1.5 ± 0.577, NPC: 20 ± 7.638, p < 0.01, **Fig. 3N**).

These findings suggest that NPC transplantation enhances axonal sprouting, neurogenesis in the SVZ, and neurite outgrowth across multiple brain regions.

### Increased vasculature density and reduced vessel leakage in mice treated with NPCs

To visualize blood vessel distribution in brain sections, coronal brain sections were stained for CD31 (**Fig. 4A, B**). 43 days after stroke induction, an ischemic border zone, characterized by hypovascularization, extended up to 300μm around the stroke core in vehicle-receiving animals (**Fig. 4C**). Vascular density, quantified here as vascular area fraction (NPC: 17%, vehicle: 9%, p < 0.001), length (NPC: 29±5.7, vehicle: 18±9, mm/mm^2^, p < 0.001), and number of branches (NPC: 651 ± 171, vehicle: 326 ± 219, mm^−2^, p < 0.001) were found significantly increased in the ischemic zone of the NPC group compared to the vehicle group (**Fig. D**). By comparison, no difference was observed between the two treatment groups in the contralateral cortex (**Suppl. Fig. 4A, B**). Interestingly, vascular area fraction and blood vessel length in the ischemic cortex recovered to similar levels as found in the contralateral intact cortex of NPC-transplanted animals.

**Fig. 4:**
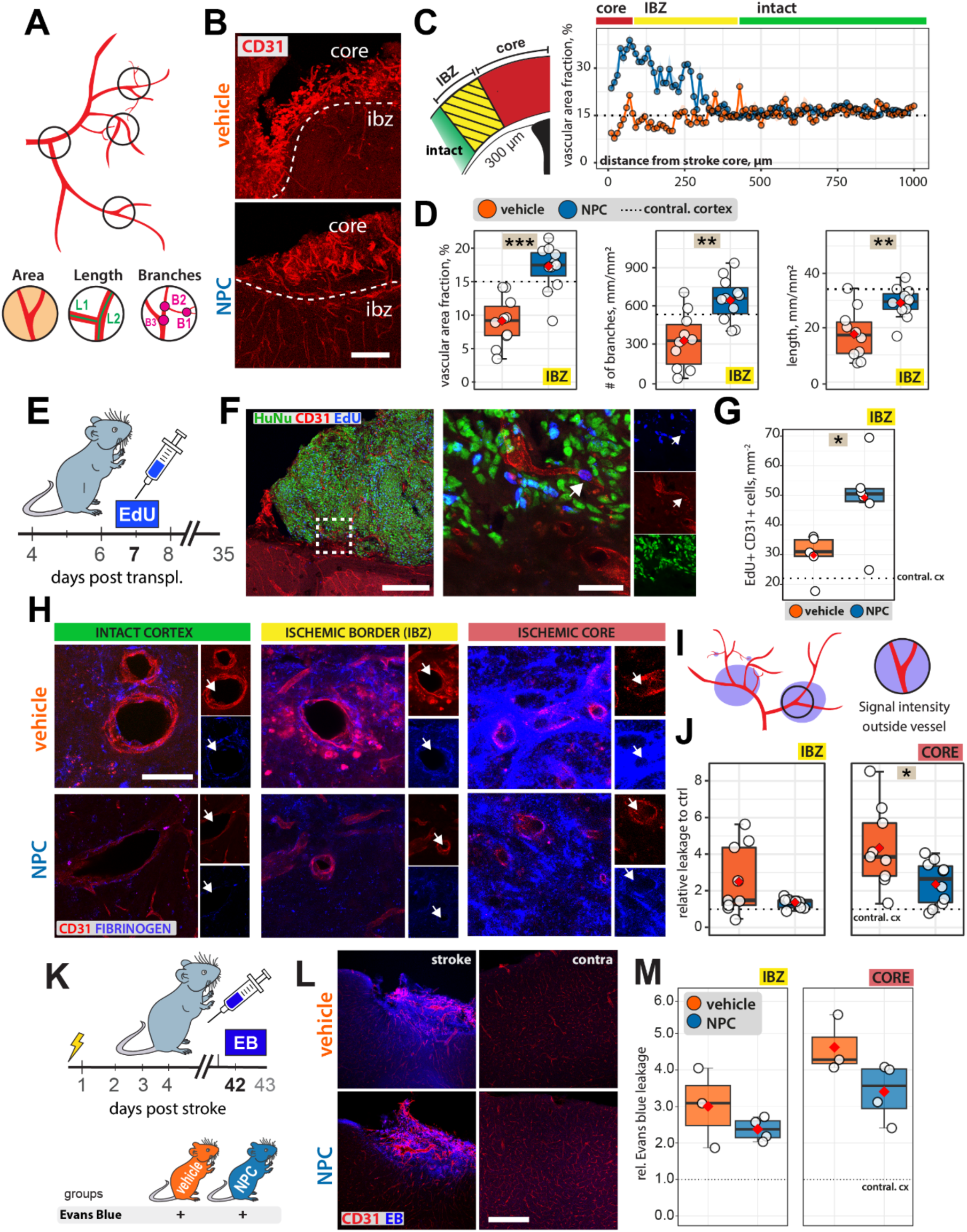
Vascular changes in ischemic stroke tissue in NPC- and vehicle-receiving animals. (A) Representative image depicting blood vessels with branch-like protrusions. (B) Immunofluorescent staining of coronal brain sections with CD31. The dotted white line indicates the ischemic border zone (IBZ). Scale bar: 100um. (C) Blood vessels were analyzed in an area up to 300μm region (yellow/green) around the stroke core (red). (D) Analysis of vasculature density (vascular area fraction, number of branches, vessel length,) in the ischemic cortex. Significances refer to comparisons between NPC and vehicle group. Dashed lines represent the contralateral hemisphere. (E) Schematic representation of EdU injection to visualize newly-formed vessels. (F) Immunofluerescent staining of coronal sections with HuNu, CD31 and EdU. Scale bar: 100um. (G) Quantification. (H) Immunofluorescent staining of coronal brain sections with CD31- (red) and fibrinogen (blue) antibodies. Segments exhibiting cross-sections of blood vessels are shown in the intact cortex, ischemic border area and ischemic core. Arrows indicate vessels. Scale bar: 10 μm. (I) Representative image depicting leaky blood vessels (J) Blood vessel leakage profiles of cortical regions in the ischemic border zone and the stroke core. (K) Schematic representation of Evans blue (EB) injection to visualize vessel leakage 24h before perfusion in control, vehicle-receiving and NPC-receiving animals. (L) Immunofluerescent staining of coronal sections with CD31 and EB. Scale bar: 100um. (M) Quantification of EB leakage relative to unaffected contralateral hemisphere in the intact cortex, the IBZ and the ischemic core. Data are shown as mean distributions where the red dot represents the mean. Boxplots indicate the 25% to 75% quartiles of the data. For boxplots: each dot in the plots represents one animal. Line graphs are plotted as mean ± sem. Significance of mean differences was assessed using an unpaired t-test (vehicle vs. NPC). In C-D, n=10 (vehicle) and n=11 (NPC) mice per group. In G, n=5 (vehicle) and n=6 (NPC). In J, n=9 mice per group; in M, n=3 (vehicle) and n=4 (NPC) per group. Asterisks indicate significance: *p < 0.05, **p < 0.01, ***p < 0.001.

To evaluate the number of newly-formed blood vessels in the ischemic border zone, the nucleotide analogue 5-ethynyl-2′-deoxyuridine (EdU) was systemically applied 7 days after cell transplantation (**Fig. 4E, F**). NPC-receiving mice showed an increase in EdU^+^ blood vessels per mm^2^ in the ischemic border zone compared to vehicle-receiving animals (NPC: 49.2 ± 14.3 mm^−2^; vehicle: 29.8 ± 7.4 mm^−2^; p < 0.05, **Fig. 4G**).

Blood vessel integrity was evaluated by utilizing vessel leakage as a parameter of vessel permeability. Immature newly formed blood vessels may cause leakage, which is linked to poor outcome after stroke^31^. To assess vessel leakage, coronal brain slices were co-stained with CD31- and fibrinogen antibodies, and sections of the ischemic border zone, ischemic core and intact contralateral cortex were analyzed in which cross-sections of blood vessels were visible (**Fig. 4H, I**). Based on the fibrinogen expression in proximity to these cross-sections, a vessel leakage profile was generated for each zone. A significant decrease in relative leakage was found in the stroke core of mice that received NPCs compared to those that had not received cells (NPC: +240%, vehicle: +435% leakage compared to contralateral side, p<0.05) (**Fig. 4J**), while no difference was observed in the IBZ of both treatment groups. As expected, there was no change in vessel leakage in the contralesional cortex of both treatment groups.

To further address blood brain barrier (BBB) integrity, animals of both treatment groups were systemically injected with Evans blue (EB), a BBB permeability marker 24 h before sacrifice (**Fig. 4K**). EB has a high affinity for serum albumin, which does not cross the intact BBB to the brain parenchyma.^32^ 43 days after stroke induction, coronal brain sections of both groups exhibited a very strong EB signal in the stroke core, and a lower signal intensity in the ischemic border zone (**Fig. 4L, M**). Compared to the contralateral cortex, both treatment groups showed a similar signal increase in both ROIs (IBZ; vehicle: +300%, NPC: +238%, stroke; vehicle: +462%, NPC: +340%, p>0.5).

Together, these data indicate that NPC transplantation is associated with increased vascular repair and maturation following stroke.

### Transplanted iPSC-derived NPCs express neuronal markers

To assess the long-term differentiation profile of transplanted NPCs, we histologically investigated the expression profile of different cell lineage specific markers. All brain sections were co-stained with HuNu antibodies or analyzed with GFP that had been transduced into the NPCs as previously described^22^ (**Fig. 5A**). Nanog, which is highly expressed in undifferentiated cells and downregulated during differentiation, was not detectable across all brain sections. A low number (12%) of transplanted NPCs still expressed the neural progenitor marker Pax6. A remarkable proportion (78%) of MAP2-positive neurons was found along with a smaller proportion (10%) of GFAP-positive astrocytes. The donut plot depicts the ratio of the individual cell types relative to all characterized cells (**Fig. 5B**).

**Fig. 5:**
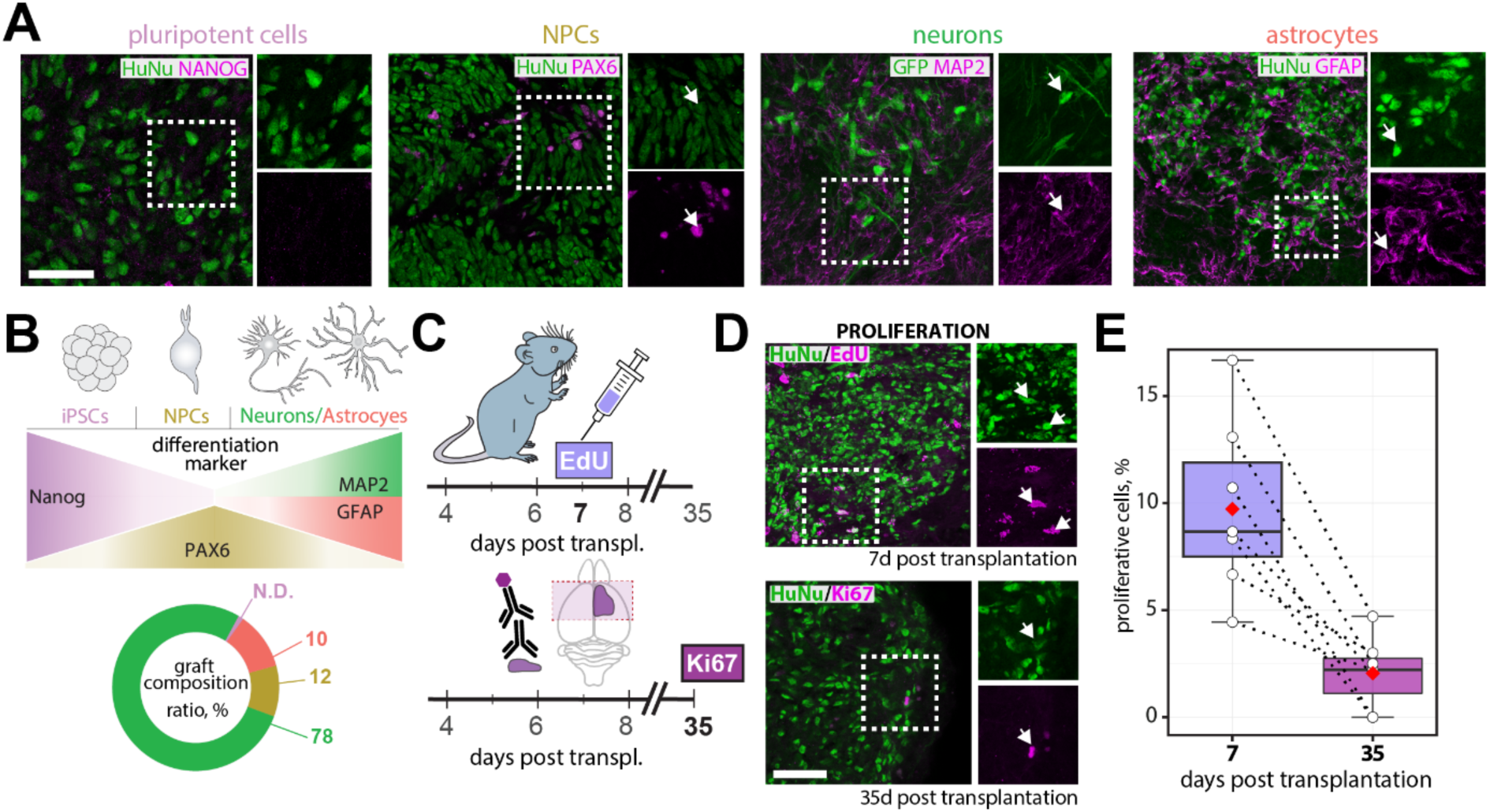
Fate of transplanted NPCs 35 days post-transplantation. (A) Representative confocal images of brain sections with Nanog, MAP2, Pax6 and GFAP, co-stained with a human-nuclei specific antibody (HuNu) or overlayed with endogenous GFP. Scale bar = 50µm (B) Quantification of Nanog+, MAP2+, Pax6+ and GFAP+ cells relative to all characterized cells. (C) Schematic representation of EdU injection to label proliferating cells 7 days post-transplantation and Ki67 antibody staining to label proliferating cells at day 35 post-transplantation. (D) Images of brain sections with EdU and Ki67, co-stained with HuNu. (E) Quantification of proliferating cells relative to all counted HuNu+ cells at day 7 (EdU) and day 35 (Ki67) post-transplantation. Scale bar = 50µm. n.d. = not detected.

We additionally co-stained with HuNu and Olig2 antibodies to determine oligodendrocyte differentiation of transplanted NPCs in the ischemic border zone and in the stroke core region (**Suppl. Fig. 5A,B**). A total of 1.1% of cells were counted double-positive for HuNu and Olig2 in the ischemic border zone, whereas only 0.54% of double-positive cells were found in the stroke core (**Suppl. Fig. 5C**).

As the *in vivo* bioluminescence imaging from the graft demonstrated a stable signal over the initial 14 days followed by a delayed increase throughout the time course of 35 days (Fig 1H), we were interested to understand the proliferation potential of transplanted NPCs at two different timepoints. The nucleotide analog 5-ethynyl-2′-deoxyuridine (EdU) was applied at day 7, while the presence of the proliferation marker Ki67 was assessed on day 35 after transplantation (**Fig. 5C, D**). A total of 10% of cells were counted double-positive for HuNu and EdU one week after transplantation, whereas only 2% of grafted cells were identified to be Ki67-positive after 35 days (**Fig. 5E**). Our findings suggest that over the initial 14-day period, the NPC count remains, despite a high proliferative potential, likely suggesting that cell proliferation and likely cell death occur at comparable rates. Subsequently, the proliferative potential of cell grafts diminishes, with a preferential differentiation into neurons.

### Transplanted NPCs improve motor function following stroke

To assess the motor function of treatment groups that underwent photothrombotic stroke, we applied previously established behavioral set-ups (rotarod assay, horizontal ladder walk, runway walk) (**Suppl**. **Fig. 6A**)^33^. Behavior assessment was conducted at multiple timepoints: pre-stroke to establish a baseline, at day 3 post-stroke to assess the severity of motor impairment, and then at regular intervals following transplantation surgery (days 7, 21, 28, 35, and 42) to monitor the recovery of motor function.

After stroke induction, a decrease in motor performance was observed in both the treatment and vehicle group in the rotarod test (−35% in NPC, −30% in vehicle) (**Suppl. Fig. 6B**). Five weeks after stroke induction, mice that received NPCs showed an improved recovery in the rotarod test compared to the vehicle group (p = 0.0021, **Suppl. Fig. 6B**).

To detect specific changes in the gait, we additionally performed a recently established automated gait analysis^33^. A customized free walking runway fitted with two mirrors allowed 3D recording of mice from lateral/side perspective and a viewpoint from below (**Fig. 6A**). The runway was exchanged with an irregular ladder rung to identify fine-motor impairments indicated by erroneous paw placement in the horizontal ladder walk test. A deep learning-based software (DeepLabCut) was used to extract and analyze pose estimation data from video footages of both tests^34^.

**Fig. 6:**
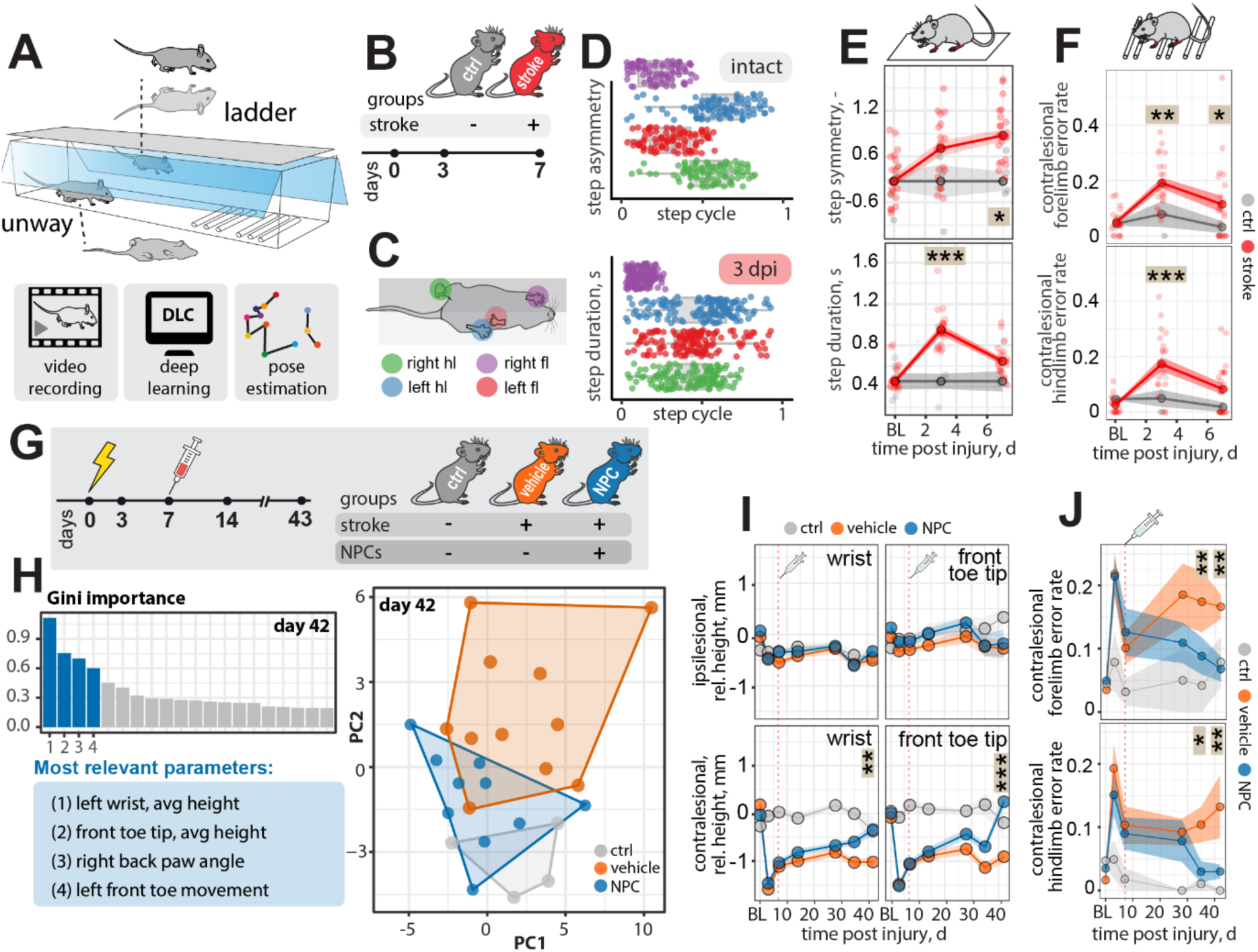
Increased recovery of motor function for mice that received NPCs. (A) Schematic representation of behavioral tests and DLC workflow used to identify and label anatomical landmarks of mice for pose estimation. (B) Experimental groups. (C, D) Footfall profile of a normalized locomotor cycle showing the stance and phase start and end of an indvidual control animal (top) and stroked animal (bottom). (E) Top: Ratio of asynchronization at baseline, 3, and 7 dpi. Bottom: Duration of a step cycle at baseline, 3 and 7 dpi. (F) Overall rate of missteps in contralesional forelimbs (top) and contralesional hindlimbs (bottom). (G) Experimental treatment groups (control, vehicle, NPC) tested for motor recovery. (H) Left: Random Forest classification of most important parameters. Right: Principal component analysis of most relevant parameters at 42 dpi. (I) Relative height of wrist and front toe tip at baseline and 3, 7, 14, 21, 28 and 35 dpi. (J) Overall rate of missteps in contralesional forelimbs (top) and contralesional hindlimbs (bottom) at baseline and 3, 7, 14, 21, 28 and 35 dpi. Missteps were detected by the back toe tips and front toe tips dipping below a defined threshold (0.5cm). Dashed line shows timepoint of transplantation. Data are shown as mean distributions where the red dot represents the mean. Line graphs are plotted as mean ± sem. Significance of mean differences was assessed using an unpaired t-test (stroke vs non-stroke) or Tukey’s HSD. In C-E, n=21 (stroke) and n=4 (no stroke) mice per group. In G-I, n=10 (vehicle), n=11 (NPC) and n=4 (ctrl) mice per group. Asterisks indicate significance: *p < 0.05, **p < 0.01, ***p < 0.001.

Individual body parts were selected to enable analysis of coordination, movement, and relative spatial positioning of the mouse joints from all three perspectives for the free walking runway and the horizontal ladder walk (**Suppl. Fig. 6C, D**). A training set of six videos proved sufficient to achieve a cross-entropy loss of <0.1% indicating a marginal predicted divergences from the actual label after 500’000 iterations (**Suppl. Fig. 6E, F)**; this indicated a highly accurate prediction for labeling all body parts. The generated neural networks were applied to all videos recorded from the runway and horizontal ladder to successfully generate a complete kinematic profile of the mouse gait (**Suppl. Fig. 7-10)**. We validated the reliability of the tracking across individual recordings (**Suppl. Fig. 7**), across days and treatment groups (**Suppl. Fig. 8**) and across different body parts (**Suppl. Fig. 9,10**), as previously established^35^.

First, injury-related gait abnormalities were assessed at baseline, 3- and 7-days (before cell therapy) following stroke (*n*=21) and data was compared to non-stroked littermates (*n*=4, **Fig. 6B**). For this, we used gait measurements that were previously identified to reliably detect acute injury^35^. The movement speed of each limb, captured from a bottom-up perspective, was used to identify individual steps (**Suppl. Fig. 10**). The footfall pattern between the front and back paws was severely altered acutely at 3dpi, here shown in a representative single-animal data (**Fig. 6C**). While the asynchronization between front/back paws was significantly increased up to 7 days after injury (p < 0.01), step duration did only differ acutely at 3dpi (p<0.01) compared to intact animals, before returning towards baseline levels (**Fig. 6D, E**). Additionally, we detected an increased error rate in the ladder rung test, affecting the contralesional hindlimbs (ipsilesional/right back paw: 0.04 ± 0.06, contralesional/left back paw: 0.18 ± 0.12, p < 0.001) and forelimbs (3dpi: ipsilesional/right front paw: 0.06 ± 0.09, contralesional/left front paw 0.18 ± 0.09, p = 0.002, 7dpi: ipsilesional/right front paw: 0.02 ± 0.04, contralesional/left front paw 0.14 ± 0.16, p = 0.027) (**Fig. 6F**).

We next applied the established behavioral set-up to the different groups (control, vehicle, NPC, **Fig. 6G**). We extracted a set of >100 parameters to examine synchronization, spatial variability, and joint angles during the spontaneous horizontal runway task, as previously described^33^. To understand the individual importance of these parameters, we applied a random forest classification at 42 dpi (**Fig. 6H, Suppl. Fig. 11A, B**). The most significant Gini impurity-based feature importance between treatment groups at 42 dpi was observed in the average heights of the left wrist and left front toe to measure long-term deficits. Interestingly, the key features identified at 42 dpi based on Gini impurity measures, showed minimal overlap from those identified at 3 dpi (**Suppl. Fig. 11B**), confirming that thorough gait assessment post-stroke is crucial for capturing the complex dynamics of functional recovery over time.

Next, we explored the most discriminative features, guided by parameters identified in prior analysis^33^. Relative height of the wrist, front toe tip, back ankle, elbow, and shoulder decreased acutely after stroke (3dpi, stroke vs ctrl: p<0.001, **Fig 6I, Suppl. Fig. 11C**). These alterations primarily affected the contralesional side but were also marginally detectable in the ipsilesional side (**Suppl. Fig. 11C**). Over the time course of 42 dpi, NPC-receiving animals experienced complete recovery, while the vehicle group showed partial but incomplete recovery in the average height of the contralesional wrist (p = 0.01), contralesional front toe tip (p<0.001) and contralesional back toe (p < 0.001, **Fig. 6I, Suppl. Fig. 11C**),

Differences in the fine-motor skill recovery of NPC and vehicle treated mice were also observed in the horizontal ladder task, The error rates were significantly higher in the contralesional forelimbs and hindlimbs of mice in the vehicle group with the most notable differences occurring at 35 and 42 days post-stroke (day 35: p = 0.005, day 42: p=0.006) and contralesional hindlimbs (day 35: p=0.031, day 42: p = 0.002, **Fig. 6J, Suppl. Fig. 12**). These findings indicate that NPC therapy leads to improved long-term functional recovery following a stroke.

To investigate whether the observed functional benefits are associated with improved peripheral muscle innervation, we examined the neuromuscular junction (NMJ) innervation status in the extensor digitorum longus (EDL) and soleus muscles (SOL) of the hindlimbs of both treatment groups (**Suppl. Fig. 13A,B**). However, we did not find any significant differences in NMJ innervation between the two groups, with more than 97.83% (EDL) and 100% (SOL) of the NMJs remaining innervated in vehicle-treated animals, and more than 97.85% (EDL) and 100% (SOL) NMJs remaining innervated in NPC-treated animals (p>0.05 for all comparisons, **Suppl. Fig. 13C**). Postsynapses of both groups show normal pretzel-like shapes and were comparable in size (vehicle_EDL_: 329±121.87, vehicle_SOL_: 373±158.1, NPC_EDL_:335.1±110.3, NPC_SOL_: 401.2±178.97, p>0.05 for all comparisons, **Suppl. Fig. 13D**). Thus, despite enhanced functional outcomes, the mechanism of recovery may not be directly reflected by changes in NMJ integrity in these muscle groups.

### Single cell molecular profiling of transplanted cell grafts

To understand the transcriptional signature and cellular identity of the grafts at cellular resolution, we performed single nucleus RNA sequencing (snRNAseq) on stroke mouse tissue containing the transplanted human cell grafts. NPCs were transplanted seven days after stroke induction, and tissue was collected and dissociated one month following transplantation for single nucleus isolation and snRNAseq, as previously described^36,37^ (**Fig. 7A,B, Suppl. Fig. 14**).

**Fig. 7:**
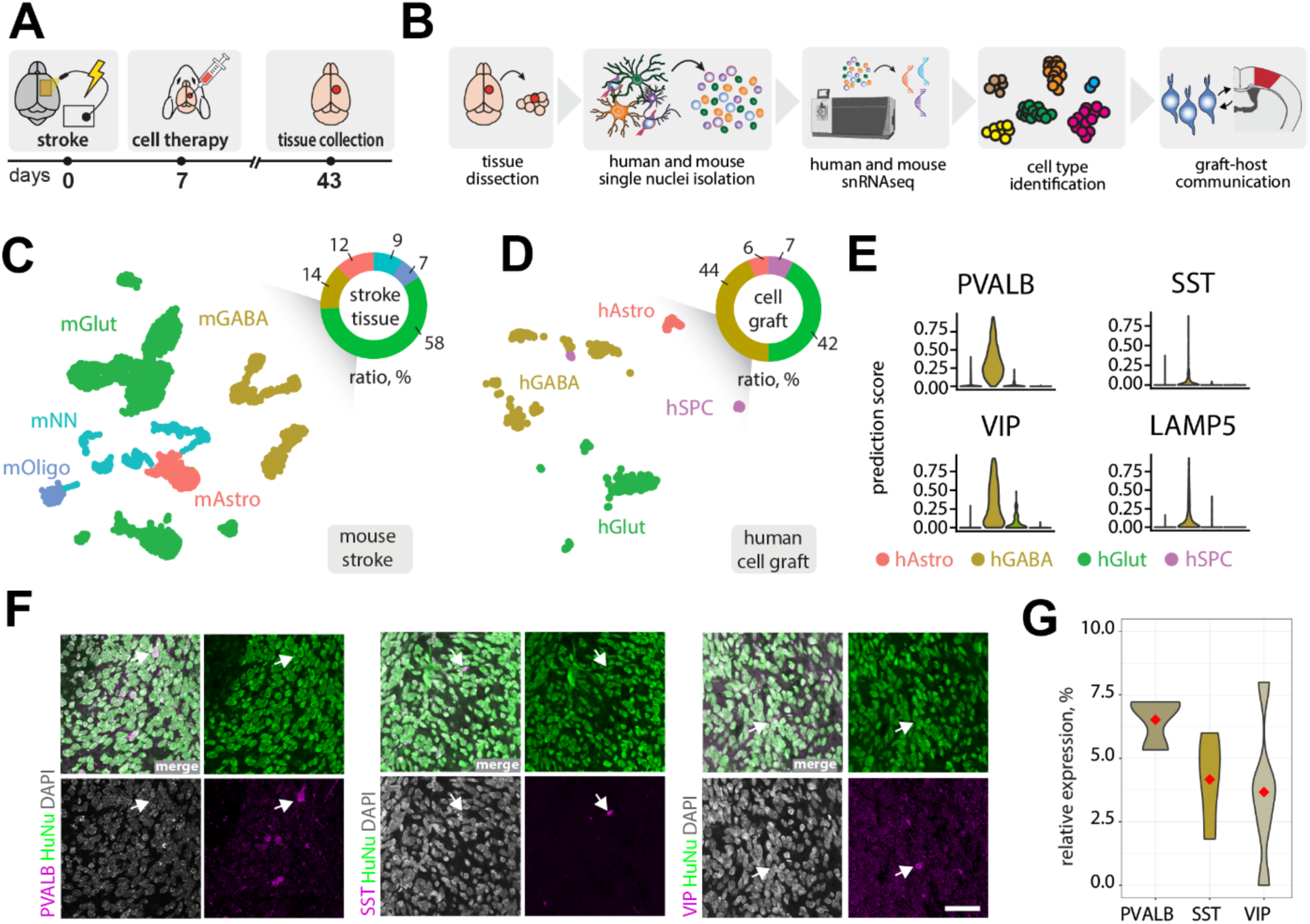
Transcriptomic profiling of single nuclei from stroked mouse host tissue and transplanted human cell grafts. (A) Experimental timeline of stroke induction, cell transplantation and sample collection. (B) Schematic representation of analysis pipeline from tissue dissociation, snRNAseq, cell type identification and graft-host communication. (C, D) UMAP and cell type identity of mouse host tissue and transplanted human cell grafts. (E) Prediction of GABAergic subtypes (in human cell graft (n = 5)). (F) Representative confocal images of brain sections with parvalbumin (PVALB), somatostatin (SST) and vasoactive intestinal peptide (VIP), co-stained with DAPI and a human-nuclei specific antibody (HuNu). Scale bar=50um. (G) Quantification of PVALB+, SST+, and VIP+ cells relative to all counted HuNu+ cells.

To define the molecular identity of the human cell graft and the surrounding stroke tissue, we used previously published human (h) and mouse (m) single cell cortical brain atlases as a reference database^38–40^ and re-classified the cell types into glutamatergic neurons (Glut), GABAergic neurons (GABA), stem and progenitor cells (SPC), oligodendrocytes (Oligo), astrocytes (Astro), and non-neural cells (NN, comprising mostly microglia, endothelial cells, fibroblasts and mural cells) (**Suppl. Fig. 15**). We also confirmed the expression of canonical cell type marker in the reference mouse brain dataset (Suppl. Fig 16). The composition of mouse and human reference datasets was very similar with Glut neurons as the most prominent cell type (human (h) Glut: 66%, mouse (m) Glut 68%), followed by GABA neurons (hGABA: 23%, mGABA 29%), astrocytes (hAstro 3%, mAstro 1%), Oligos (hOligo: 4%, mOligo 1%), non-neural cells (hNN: 2%, mNN: 1%). The human dataset also consisted of 2% of stem and progenitor cells (containing PAX6-expressing NSC/NPCs and oligodendrocyte progenitor cells), which were particularly useful to identify and map non-differentiated NPC cell grafts (**Suppl. Fig. 15**).

In total 83’058 nuclei from five pooled mouse stroke brains were mapped after pre-processing with an overall confidence of >99% (mAstro = 71%, mGABA = 100%, mGlut = 100%, mNN = 89%, mOligo = 97%) to the mouse cortex reference atlas (**Suppl. Fig. 16, Suppl. Fig. 17**), and 1149 human graft nuclei were mapped with an overall average confidence of >90% (hAstro = 68%, hGABA = 84%, hGlut = 94%, hSPC = 62%) to the human cortex reference atlas (**Suppl. Fig. 17C,D**).

Compared to the mouse reference data the number of non-neural nuclei strikingly increased in the stroked mouse tissue (**Fig. 7C**). Interestingly, while the relative population of glutamatergic cells was only slightly reduced after stroke (intact mGlut 68%, stroked mGlut: 58%), the number of GABAergic cells was halved (intact mGABA: 29%, stroked mGABA: 14%), as previously described by other studies linking GABAergic interneurons to stroke recovery^41–43^. As expected, we observed within each analysed cell type differentially expressed genes (DEGs) with most prominent changes in mGABA and mGlut neurons (**Suppl. Fig. 18, Suppl. Table 2**), similar to previous studies^44,45^. Gene Set Enrichment Analysis (GSEA) of these DEGs revealed distinct biological processes associated with stroke-induced transcriptional changes in each cell type that are primarily downreulated, with astrocytes showing decrease in metabolic pathways, GABAergic neurons exhibiting alterations in synaptic signaling, and non-neuronal cells and oligodendrocytes showing changes related to energy metabolism (**Suppl. Fig. 19, Suppl. Table 3**).

Human nuclei from transplanted NPCs showed 94% mature cell characteristics, differentiating into astrocytes (6%), as well as GABAergic (44%) and glutamatergic neurons (42%). Only 7% of transplanted cells showed characteristics of stem and progenitor cells (**Fig. 7D**), similar to our observations from immunohistochemical phenotyping in stroke sections (**Fig. 5**). The majority of grafts (44%) were associated with the GABAergic phenotype and more detailed analysis revealed characteristics of parvalbumin (PVALB), somatostatin (SST), vasoactive intestinal peptide (VIP), and lysosomal-associated membrane protein 5 (LAMP5) inhibitory interneurons (**Fig. 7E, Suppl. Fig. 20**). Graft expression of PVALB, SST and VIP was further confirmed by immunohistochemical staining (**Fig. 7F,G**). These findings suggest that transplanted cell grafts can acquire a GABAergic-like phenotype, potentially replacing the GABAergic neurons lost in stroke-damaged brains.

### Decoding cell-graft communication at single cell resolution

We observed that the proportion of GABAergic neurons in the mouse stroke tissue was considerably reduced compared to intact mouse brain; and that half of the transplanted NPCs acquired a GABAergic-like phenotype after stroke. We hypothesized that GABAergic grafts interact with damaged host tissue to initiate tissue repair processes. Using CellChat^46,47^, we quantitatively analyzed interactions of ligands and receptors between graft and host single cells (**Fig. 8A**). First, we identified human orthologues for mouse data and explored the number and weight of interactions between graft and host (**Fig. 8B,C, Suppl. Fig. 21**). As expected, most interactions occurred between host mGABA and mGlut cells via both autocrine and paracrine signaling (**Fig. 8B**) but we also observed interactions of all graft cells with all host cell types, most prominent interactions were between hGlut with mGABA and mGlut, as well as hGABA with mGABA, mOligo and mAstro. Interestingly, the number of predicted incoming and outgoing intercellular interactions was highest for hGABA cells among the graft cells (**Fig. 8D**).

**Fig. 8:**
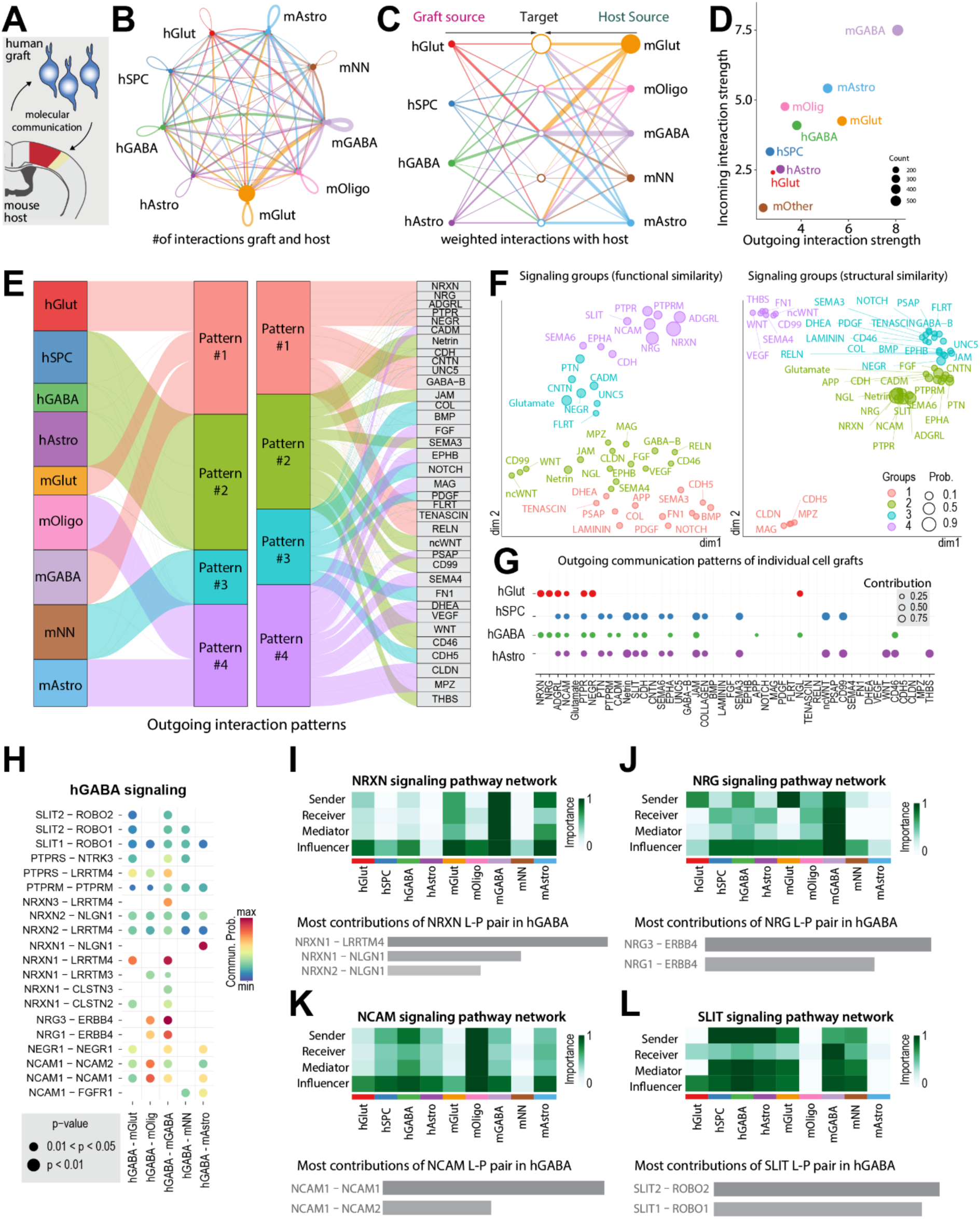
Molecular crosstalk between cell graft and host. (A) Schematic of graft and host interaction analysis (B) Total number of cell-cell interactions between individual cell types of graft and host (C) Hierarchy plot of weighted interaction between graft and host source cell types to host target cell types. Circle sizes are proportional to the number of cells in each cell type and edge width represents the communication probability in B and C. (D) Projecting incoming (y axis) and outgoing (x axis) interaction strength of individual graft and host cell types in a 2D space. Each dot represents the communication network of a cell type. Dot size is proportional to the overall number of cells in each cell group. (E) Riverplot of four outgoing interaction patterns for each cell type. The thickness of the flow indicates the contribution of the cell type to each pattern. (F) Projecting signaling groups onto a 2D space according to their functional (left) and structural (right) similarity. Each dot represents the communication network of one signaling pathway. Dot size is proportional to the overall communication probability. Different colors represent different groups/patterns of signaling pathways. (G) Outgoing communication patterns of individual graft cell types (H) Major ligand-receptor pairs that contribute to the signaling sending from human (hGABA) cells to host cell types from stroked tissue. The dot color and size represent the calculated communication probability and p-values. p-values are computed from one-sided permutation test. (I) Heatmap and barplot show the relative importance of each cell type based on the computed four network centrality measures (sender, receiver, mediator, or influencer) of NRXN signaling, (J) NRG signaling, (K) NCAM signaling, (L) SLIT signaling.

In addition to intercellular communications, we explored communication patterns that connect cell groups with signaling pathways in the context of outgoing signaling and incoming signaling (**Fig. 8E, Suppl. Fig. 22**). We categorized the signaling in four signaling groups that could be separated by functional and structural similarity (**Fig. 8E, F**). We identify that a large portion of outgoing signaling from hGlut and hGABA neurons fits into pattern #1 and pattern #2, representing pathways such as neurexin (NRXN), neuregulin (NRG), neural cell adhesion molecule (NCAM), Netrin and semaphorin3 (SEMA3) (**Fig. 8E-G, Suppl. Fig. 23**). These pathways have been associated with neuronal regeneration, synapse plasticity, dendritic arborization and axon guidance^48–51^.

Since the host cells that were lost after stroke were disproportionally mGABA neurons, we explored whether graft-derived hGABA neurons may contribute to outgoing signaling to individual host mouse cells. The hGABA grafts signaled to all host cell types. Some signaling pathways were specific to individual graft cell types e.g. hGABA to mAstro via NRXN1 – neuroligin 1 (NLGN1), while other signaling mechanisms may involve several host cell types e.g., NRG1/3 to erb-b2 receptor tyrosine kinase 4 (ERBB4) signaling occurs with both mOlig and mGABA cells (**Fig. 8H, Suppl. Fig. 24, 25**). Using computed network centrality measures, we explored the most significant ligand-receptor pairs: NRXN, NRG, NCAM and SLIT and the role of graft cells in the signaling as dominant senders, receiver, mediator, or influencers for the intercellular communication (**Fig. 8I-L**). While hGABA cells preferentially acted as influencers in NRXN signaling, they were the most prominent sender between graft cells in NRG, NCAM and SLIT signaling. The relative contribution of each ligand-receptor pair in hGABA cells to the overall communication network in NRXN, NRG, NCAM and SLIT signaling pathways are: NRXN1 – LRRTM4 with mGlut and mGABA cells (**Fig 8I, Suppl. Fig. 25**), NRG3 - ERBB4 with mOlig and mGABA cells (**Fig 8J, Suppl. Fig. 25**), NCAM1 – NCAM1 with most host cell types (**Fig 8K, Suppl. Fig. 25**), and SLIT2 – roundabout guidance receptor 2 (ROBO2) with mGlut and mGaba cells (**Fig. 8L, Suppl. Fig. 25**). Notably, hGlut grafts were also predicted to contribute to outgoing communication and act as influencers (e.g. exerting a regulatory effect within a network), particularly within NRXN, NRG and NCAM signaling **(Fig. 8G**, **Fig. 8I-L**). However, their overall interactions appeared more limited compared to hGABA grafts, with fewer connections in NRXN, NRG, and NCAM signaling and no predicted engagement in SLIT signaling (**Suppl. Fig. 25**). Major pathways that were predicted to contribute to outgoing communication of hGlut cells included neuronal growth regulator (NEGR), adhesion G protein-coupled receptor-like (ADGRL), and protein tyrosine phosphatase receptor (PTPR) signaling pathways (**Suppl Fig 26**). Among the top outgoing interactions from hGlut cells, the TENM3–ADGRL3 and NEGR1–NEGR1 ligand–partner pairs suggest graft-host interactions that may facilitate synaptic communication and adhesion to host brain circuits (**Suppl. Fig 26**).

As a proof-of-concept, we performed a Boyden chamber assay to assess whether our NPCs migrate in response to Slit2 (**Suppl. Fig. 27)**. Cells were seeded in transwell inserts and exposed to different concentrations of mouse Slit2 protein (0, 100, 200 ng/ml). After 24 hours, a significant increase in NPC migration was observed toward 100 and 200 ng/ml Slit2, whereas minimal migration was detected with no Slit2 (0 ng/ml). This result suggests that moderate levels of Slit2 may promote NPC chemotaxis and support a role for Slit2 signaling in the graft’s migratory response. Taken together, these findings suggest molecular communication between cell grafts and the surrounding stroke-injured tissue through regeneration-associated pathways.

## Discussion

Here, we have demonstrated that transplanted NPCs engraft and survive for at least five weeks in stroke-infarcted tissue. Mice that received the cell grafts exhibited reduced inflammatory response and increased post-ischemic angiogenesis in the ischemic cortex, along with improved vessel integrity in the stroke core, and increased recovery of motor function. We have provided molecular definitions of graft-derived cells and propose preferential differentiation into glutamatergic and GABAergic neurons. Graft-derived GABAergic neurons communicate with stroke-injured host tissue likely through NRXN, NCAM, SLIT and NRG signaling pathways, which are associated with neuronal regeneration, synaptic plasticity, dendritic arborization and axon guidance.

The here applied NPCs were generated using a previously established protocol, which is xeno-free and chemically defined^22^. This protocol does not require selection or purification and can thus be adapted to GMP-grade for clinical applications. Additionally, we have previously demonstrated that this cell source is compatible with safety switch systems, such as the RNA switch technology, to selectively exclude pluripotent cells before transplantation, further improving their safety profile^22^. Building on prior findings, which demonstrate that neural grafts remaining viable beyond the initial 12 days post-transplantation often endure for at least eleven months within the brain^52^, we suggest that our grafted NPCs might similarly sustain prolonged survival beyond five weeks. In addition to graft safety and quality considerations, the timing of transplantation is critical for achieving robust donor cell survival and engraftment. Delivering NPCs at day 7 post-stroke, when excitotoxicity and edema have partially subsided and inflammatory signals are shifting, has been commonly reported in the literature^53–55^. In contrast, we have found that earlier transplantation (24 hours post-stroke) resulted in significant lower graft survival within the first days after transplantation^56^. Consequently, synchronizing the intervention with the changing conditions of the host brain appears essential for maximizing graft survival and therapeutic efficacy. Clinically, there is also significant interest in cell therapies for patients in the chronic stroke phase, a stage where the majority of stroke survivors currently reside and where treatment options remain limited. Previous research suggests that there is a long-lasting critical period of enhanced neuroplasticity post-stroke, enabling improvements in function/structure even long after the initial injury.^57,58^ A recent preclinical study further supports this notion, showing that hMSC therapy can correct stroke-induced neural circuit disruptions even at chronic stages by reopening a period of adaptive plasticity.^62^ This finding suggests a broad potential for stem cell therapies in addressing neurological conditions characterized by network hyperexcitability. Clinical trials have demonstrated both the safety and potential therapeutic benefits of stem cell interventions at chronic stages of stroke.^59–61^ For instance, significant motor and neurological recovery improvements have been observed in patients who received stem cell treatments >90 days after severe middle cerebral artery territory infarct, highlighting the persistence of a therapeutic window far beyond acute injury.^14^ One issue with transplantation in chronic phases is that injections can be very harmful to adjacent brain tissue where important recovery processes may already be underway.^63^ Therefore, innovative strategies have been developed, including combining cell therapy with biopolymer scaffolds composed of extracellular matrix molecules such as collagen and hyaluronic acid, providing a protective environment for transplanted cells and minimize disruption to surrounding brain tissue^64^

Neuroinflammation strongly contributes to the hostile post-stroke environment, and, among other aspects, it triggers glial cell activation. This process plays a dual role: astrogliosis is essential for repair processes and glial scarring, and microglial activation is associated with proinflammatory response and tissue injury^65^. Significant downregulation of GFAP expression after stem cell-based therapy for acute ischemic stroke has been reported in studies using MSCs^66,67^. Compared to NPCs, MSCs are unable to differentiate into innate brain cells but may be more prone in releasing factors that dampen astrocyte reactivity. For decades, the prevailing dogma dictated that glial scarring is a key inhibitor of CNS regeneration and a barrier to axonal outgrowth by inhibiting axonal regeneration^68–70^. However, more recent evidence suggests that this scar strongly supports axon regeneration in spinal cord injury, and axons are capable of regrowing through dense astrocytic scar tissue^71^. Indeed, we observed neurite extension from the graft to various brain regions, suggesting that transplanted cells may navigate or even modify the glial scar. Vice versa, the glial scar might contain structural and chemical cues that are critical for functional integration of cell grafts^72^. Further investigations will be needed to determine exactly how transplanted NPCs interact with resident glial elements to promote neurite outgrowth. While still highly debated, our study supports the notion that glial scarring may actually provide a supportive scaffold for regeneration under certain conditions, rather than being purely inhibitory.

Although acute inflammation is necessary to induce tissue repair, collateral damage by a persistent inflammatory response exacerbates ischemic damage and is associated with poor prognosis^73,74^. NPCs have previously been reported to release neurotrophic factors such as glial-derived neurotrophic factor (GDNF), brain-derived neurotrophic factor (BDNF) and nerve growth factor (NGF) that inhibit down-stream effects of inflammation^75–77^. In addition, studies have confirmed decreased microglial activation through reduction of pro-inflammatory cytokines after intraparenchymal transplantation of hiPSC-NPC^30,78^.

Post-stroke inflammation further leads to BBB breakdown, which has been linked to poor outcome after stroke, as tissue becomes inefficiently supplied with oxygen and glucose^31^. Previous preclinical approaches have focused on promoting angiogenesis through the application of vascular growth factors, such as vascular endothelial growth factor (VEGF). Yet, VEGF is known to initiate a series of detrimental events, such as increased vascular permeability, BBB breakdown, and an elevated risk of bleeding from immature, unstable vessels (hemorrhagic transformation)^79,80^. We report that NPC-treatment leads to significantly increased blood vessel density in the ischemic core. We hypothesize that NPC grafts secrete angiogenic factors that promote vascular repair, vessel maturation and successfully support tissue regeneration while reducing the risk of vascular leakage and bleeding. Previous reports have shown that cortical infarcts produce a temporally restricted window of heightened vascular structural plasticity, most prominently within the first two weeks after stroke^81^. During this interval, newly formed capillary connections can help reestablish blood flow in peri-infarct tissue, thereby alleviating local ischemia, supporting synaptic remodeling, and ultimately enhancing functional recovery. Notably, we transplanted our NPCs on day 7 post-stroke, aligning with this critical plasticity window. Therefore, the pro-angiogenic cues from the grafted cells likely synergized with the endogenous remodeling processes to promote vascular repair.

Cerebral ischemia induces neurogenesis in the subventricular zone (SVZ), followed by the migration of these newly generated neuroblasts to the injury site, where they differentiate into functional neurons^82,83^. However, this self-repair process is inherently limited and has therefore become a prime target for therapeutic interventions. Cell therapy has been shown to assist in this process through paracrine mechanisms^7,11,24^. In line with these previous reports, we demonstrated that NPC transplantation increased endogenous neurogenesis in the SVZ. Although newly generated neurons appear to represent only a small fraction of all SVZ-derived cells responding to stroke, they have been linked to improved functional outcomes by contributing to synaptic and vascular repair and by providing trophic support^83,84^. Indeed, ablation or reduction of SVZ neural stem cells, which diminishes this neuronal output, leads to poorer recovery. Hence, even the relatively modest population of newborn neurons migrating from the SVZ plays an important role in promoting repair mechanisms.

The percentage of dividing grafted NPCs considerably decreased from day 7 post-transplantation over a 3-week period. It is well-established that during development, asymmetrical division of NPCs leads to self-renewing daughter cell and a post-mitotic neural cell subtype prior to terminal differentiation^85^. Besides, it is important to note that the increased bioluminescent signal is not associated with any aberrant cell transformation or growth, which we already documented earlier^22^. We therefore suggest that the cell grafts undergo controlled limited cycles of proliferation before maturation into neurons and glial cells. This hypothesis is supported by the absence of undifferentiated cells and the scarce presence of Pax6^+^ NPCs at day 35 post-transplantation, which is similar to previous observations^10^. The fate and function of these residual progenitor cells remain unknown, and it should be further analyzed in longer-term studies.

We found that half of the transplanted NPCs differentiated into neurons with a GABAergic phenotype. Previous studies have documented similar ratios of terminal differentiation of NSCs/NPCs into neurons and glia^86–92^. This is especially relevant given that the GABAergic cell population experienced a reduction of almost 50% after the ischemic insult. While glutamatergic neuron replacement has received widespread attention in this field of research^9,93^, there has been little emphasis on addressing ischemia-induced neuronal depletion and replacement specifically in GABAergic interneurons. Indeed, restoration of GABAergic neurotransmission can potentially decrease the damage produced by excitotoxicity in the stroke acute phase^94^. Optogenetic activation of inhibitory neurons at 24h post-injury results in better stroke outcomes^95^. A more detailed analysis of GABAergic cells revealed characteristics of PVALB, VIP, SST and LAMP5 inhibitory interneurons. Optogenetic investigations of PVALB interneurons have shown that, while their excitability recovers after reperfusion, GABAergic synaptic activity remains impaired for more extended periods^96^, which highlights the complexity of the inhibitory neural network. Indeed, since the contralesional side exerts an interhemispheric inhibition^97^, optogenetic inactivation of PVALB neurons in the contralateral hemisphere side reduces infarct volumes and improves motor function in ischemic rats^98^. Chemogenetic activation of VIP neurons, whose function is disrupted after stroke, leads to enhanced sensory responses in the stroke-affected cortex through disinhibition thereby improving functional recovery^99^. Further, overexpression of Lypd6 in SST interneurons reduces PVALB activity, and therefore, increases plasticity and visual acuity through disinhibition of excitatory neurons^100^. Cell-cell interaction analysis further revealed that GABAergic interneurons likely mediate endogenous stroke recovery through NRXN, NCAM, SLIT and NRG signaling pathways in our model. NRG has been reported to act in GABAergic interneurons activating neuron firing in response to changes in excitatory activity via its ligand ERBB4^101,102^. NRG-ERBB4 signaling appears to attenuate neuronal cell damage under oxygen glucose deprivation and ameliorates pathological increases in BBB permeability after cortical impact^101,102^. In addition to the protection against neuronal death in microinfarcts, SLIT signaling has been linked to the inhibition of neuroinflammation^103,104^. Notably, in our recent single-cell transcriptomic atlas of permanent focal ischemia at one month post-injury, we found that SLIT signaling pathways were upregulated in the peri-infarct tissue, further highlighting the importance of SLIT in supporting tissue remodeling and curbing neuroinflammation after stroke^45^. Given that our transplanted NPCs communicate with the host brain through similar pathways, it is possible that cell therapy may further support these endogenous reparative mechanisms. Neurexins, on the other hand, are widely thought to promote synapse formation; combination of NRXN-1b and NRG-1 has been demonstrated to induce synaptic differentiation and neurotransmitter exchange^105,106^.

This set of studies has several limitations. The use of immunodeficient Rag2^−/−^ mice, rather than fully immunocompetent wildtype animals, may limit the direct clinical translation of our data, as immune responses to grafted cells could affect graft survival and functional outcomes. Future work will pursue the use of engineered hiPSC-NPC lines that evade immune rejection and can serve as allogenic therapies for stroke^107^. Second, while we have demonstrated that grafted iPSC-derived NPCs can differentiate into mature neurons with long-range axonal projections, we did not directly assess their functional integration. Other studies have demonstrated that grafted human iPSC-derived cortical neurons send axonal projections across hemispheres in stroke-injured rats, form excitatory glutamatergic synapses on host cortical neurons, and become partially integrated into the host sensorimotor circuit^9^. Finally, while our data support the efficacy of NPC transplantation, further mechanistic validation remains necessary. Similar studies have used diphtheria toxin (DT) to selectively ablate human cells in rodent brains, demonstrating a feasible strategy to test whether therapeutic effects are directly attributable to transplanted cells, secreted trophic factors, or both^10^. Future work could employ similar approaches to dissect the precise mechanisms by which NPCs promote stroke recovery.

In summary, our work establishes a preclinically effective iPSC-derived neural cell source for stroke recovery and decodes the underlying molecular graft-host communications. These findings could have relevance not only for stroke therapy, but also for treating a broader range of acute neurological injuries that involve neuronal loss.

## Materials and Methods

### Experimental Design

This study was designed to assess the efficacy and regenerative capacities of a human hiPSC-NPC therapy for stroke in immunodeficient *Rag2*^−/−^ mice. We hypothesized that the intraparenchymal transplantation of NPCs seven days after stroke could lead to (1) tissue repair responses and (2) an improved recovery in motor function. We chose the time point of transplantation based on previous findings^56^. To test this, mice were allocated into one of three treatment groups: (1) the first group (*n*=11) received a photothrombotic stroke and underwent NPC transplantation, (2) the second group (*n*=11) received a photothrombotic stroke and sham transplantation, and (3) the third group (*n*=4) received sham surgery and sham transplantation, serving as negative control. Photothrombotic stroke induction was validated by performing Laser-Doppler Imaging (LDI) 30 min after stroke onset, and additionally before perfusion 43 days post-injury. To determine efficacy of the NPC treatment, several parameters were considered as read-outs. Over the time course of 5 weeks, motor function was assessed for all treatment groups at different time points by performing several behavioral tests (rotarod test, horizontal ladder walk, horizontal runway walk). Video recordings of the horizontal ladder- and runway walk were analyzed with DeepLabCut (DLC) to create comprehensive 3D locomotor profiles. To track survival and migration of transplanted NPCs, mice were subjected to bioluminescence imaging (BLI) on the day of NPC transplantation, four days later, and on a weekly basis thereafter. After sacrifice, histological analysis was performed to characterize grafted cells and evaluate expression levels of cell markers that have key functions in stroke pathology. In addition, stroke size and graft size analysis were performed. All animals are presented in the study; no statistical outliers were excluded.

Both sexes where used and analyzed for this study, our statistical analyses did not reveal any significant sex-related differences in the measured parameters.

### Animals

All animal experiments were performed at the Laboratory Animal Services Center (LASC) in Schlieren according to the local guidelines for animal experiments and were approved by the Veterinary Office in Zurich, Switzerland. In total, 28 adult male and female genetically immunodeficient *Rag2*^−/−^ mice (weight range: 20-30g) were employed for this study. Mice were kept at the LASC in top-filter laboratory cages under OHB conditions in a temperature and humidity-controlled room with a constant 12/12h light/dark cycle (light on from 6:00 a.m. until 6:00 p.m.). All mice were standard housed in groups of at least two and maximal four per cage with ad libitum access to standard diet food pellets and water.

### hiPSC-NPC culture

For a cell-based stroke therapy, our group has established a protocol to differentiate human iPSCs to NPCs under chemically defined, xeno-free conditions^22^. On day –2, 80,000 iPSCs per well were plated on vitronectin-coated 12-well plates in StemMACS iPS-Brew XF medium supplemented with 2 μM Thiazovivin and incubated overnight. The next day, the medium was replaced with fresh medium. On day 0, neural differentiation was initiated using a neural induction medium (50% DMEM/F12, 50% Neurobasal medium, 1 × N2-supplement, 1 × B27-supplement, 1 × Glutamax, 10 ng/ml hLIF, 4 µM CHIR99021, 3 µM SB431542, 2 µM Dorsomorphin, 0.1 µM Compound E), with a medium change on the following day. On day 2, the medium was switched to Neural Induction Medium 2 (50% DMEM/F12, 50% Neurobasal medium, 1 × N2-supplement, 1 × B27-supplement, 1 × Glutamax, 10 ng/ml hLIF, 4 µM CHIR99021, 3 µM SB431542, 0.1 µM Compound E) for four additional days, and replaced daily. On day 6, cells were transferred to pLO/L521-coated plates in Neural Stem Cell Maintenance Medium (50% DMEM/F12, 50% Neurobasal medium, 1 × N2-supplement, 1 × B27-supplement, 1 × Glutamax, 10 ng/ml hLIF, 4 µM CHIR99021, 3 µM SB431542). NSMM was changed daily, and cells were passaged at 80–100% confluency. From passage two onward, 5 ng/ml FGF2 were added and during the first six passages, 2 μM Thiazovivin was added after splitting.

Gene- and protein expression analysis showed stable upregulation of NPC markers (Pax6, Sox1, and Nestin) and downregulation of iPSC markers (Oct4) over 15 passages. Starting with 10^6^ iPSCs resulted in an estimated 10^18^ NPCs after 15 passages. Neural progenitors were tested for potential neural differentiation. Withdrawal of growth factors (FGF2, hLIF) and small molecules (CHIR99021, SB431542) resulted in spontaneous differentiation into β-III-Tubulin- and MAP2-positive neurons as well as S100B-positive astrocytes. Prior to transplantation, NPCs were transduced with a lentiviral vector comprising a bioluminescent (red firefly luciferase) reporter and a fluorescent (eGFP) reporter.

### Cell migration assay

NPCs migration was analysed in vitro using the Boyden chamber assay. Millicell® Hanging Cell Culture Inserts with 8 μm pores (Merck) were placed on a 24-well plate and coated for 2h using Poly-L-Ornithin (PLO, Sigma Aldrich), followed by 2h incubation with Biolaminin 521 LN (L521, Biolaminin LN). 0.2mL of NSMM+SM containing dissociated cells (5 ×10*^5* cells per well) was added onto the insert. In the bottom chamber, 0.9mL of NSMM+SM with 0ng/ml, 100ng/ml or 200ng/ml Slit2 (R&D) was added. 24h after seeding, media was removed from the bottom chamber and cells attached to the upper side of the membrane filter were scraped off with a moistened cotton swab. Cells attached to the bottom side of the membrane filter were stained with DAPI and imaged with DMi8 microscope.

### Photothrombotic stroke induction

Photothrombotic stroke was induced as previously described^32,108,109^. Anesthesia was induced with 4% isoflurane vaporized in oxygen. Once the breathing rate of the mice reached a frequency of approximately 50 breaths per minute, indicating a deep state of anesthesia, the mice were removed from the induction chamber and were placed head-first into a *Davids Kopf Instruments* stereotactic frame for surgery. A self-fabricated face mask consisting of a rubber tube with a small cut-in vent enabled continuous anesthesia with a lower concentration of isoflurane (1-2%). Body temperature was kept constant at 36-37 °C using a heating pad placed underneath the surgical setup. Toe pinch reflex initiation was attempted by pinching the animal’s webspace area between the thumb and the second proximate finger to ensure deep anesthesia. After the animal showed no sign of a reflex, eye lubricant (Vitamin A, Bausch&Lomb) was smeared onto the eyes to prevent dryness throughout the surgical procedure. The animal’s head was shaved with an electric razor from the neck up to the beginning of the snout. *Emla™* Creme 5% (consisting of lidocaine and prilocaine) was then spread onto the scalp with a Q-tip and put into the ears. Ear bars were then gently inserted into the lidocaine-saturated ears underneath the ear bone to fixate the skull. An approximately 1 cm incision was made along the longitudinal fissure, and the skin was retracted by hand far enough to expose Lambda and Bregma. The fascia was removed from the skull with a Q-tip, and the periosteum of the lower right hemisphere was scraped off with a surgical scalpel. Using an *Olympus SZ61* Surgery microscope, bregma was located. With the help of a *WPI UMP3T-1* stereotactic coordinate system, the exact locus of stroke induction [-2.5mm to + 2.5mm medial/lateral and 0 mm to + 3 mm from Bregma] was identified and marked with a pen. Rose Bengal dissolved in sterile 0.9% NaCl solution[15mg/ml] was injected intraperitoneally [10μl/g bodyweight] 5 min prior to illumination with a standard insulin needle to ensure systemic circulation. An Olympus KL1500LCD (‘’4’’) cold light source illumination lamp [150W, 3000K] with a surface area of 4mm x 3mm was positioned onto the previously marked locus, and illumination ensued for 10 minutes. Once illumination ended, the mouse was placed into an empty cage. 30 minutes post-surgery, the mouse’s skull was imaged using LDI to confirm stroke induction. After stroke verification, the incision was sutured using a Braun surgical suture kit. Betadine® was applied to the suture, and the mouse was returned to an empty cage, followed by standard post-op care.

### NPC transplantation

Transplantation of NPCs was conducted as described before^110^. Operation procedures remained the same with photothrombotic stroke surgery until the point of illumination. After locating bregma, the stereotactic device was adjusted to the coordinates of the injection site along with the depth coordinates [AP: + 0.5 / MV: + 1.5 / DV: - 0.6 (mm relative to bregma)]. A small hole of ca. 0.8 mm in diameter was drilled into the skull until the cortical bone was penetrated. A 10 μl Hamilton syringe attached to a 30G needle filled with 2.5 μl of cell suspension (10^5^ cells /μl) was mounted to the stereotactic device and placed above the target site. The needle was guided to the calculated depth at a slow, steady rate. An additional 0.05 mm depth penetration was completed to create a pocket for the cells to avoid spillover of the cell suspension. Once the needle was retracted back to the calculated depth coordinate, cells were injected at a constant rate (2nl/s) into the brain parenchyma. The needle was left in place for 5 min after cell injection to allow the suspension to settle before slow withdrawal. Histoacryl^®^ was applied to seal the cavity. Once the Histroacryl hardened, the skin was sutured, and the animal was returned to an empty cage, followed by standard post-op care.

### Bioluminescent imaging (BLI)

Animals were imaged using the IVIS^®^ Lumina III In Vivo Imaging System. The entire head region was shaved with an electric razor to achieve optimal imaging. Mice were injected intraperitoneally with diluted D-luciferin dissolved in PBS (300 mg/kg body weight, sterile-filtered through a 0.22 μl syringe filter). Images were acquired 10, 15, 20, and 25 minutes after substrate injection. Imaging data attained by IVIS was analyzed with the *Living Image* software (V 4.7.3). A region of interest (ROI) was set (height, width = 1.8cm) and placed directly over the color-coded signal between the ears and nose of each animal. Two further ROIs were fixated at the posterior part of the animal’s body and at an additional randomly selected region on the image proximal to the body; the former to quantify background ROI, the latter to quantify image noise. Signal was quantified using total photon flux (photons per second). Data was recorded in Microsoft Excel and plotted with R software.

### Laser Doppler imaging (LDI)

The surgical set-up was sanitized, and the mice were placed in a stereotactic frame. The skull was exposed through a midline skin incision. Mice were subjected to LDI (*Moors Instruments, MOORLDI2-IR)* immediately after stroke induction. LDI data was exported and quantified in terms of flux in the ROI using Fiji (ImageJ) and analyzed with R software.

### Isolation of nuclei from frozen brains

Single nuclei were isolated as previously described^37^. Briefly, cortical brain tissue was removed and flash frozen using liquid nitrogen. Cortices were then homogenized using a Dounce homogenizer in lysis buffer (10mM Tris-HCl, ph 7.5, 10mM NaCl, 3mM MgCl2, and 0.1% Nonidet P40 in nucleuase-free water). Homogenized tissue was incubated for 15 minutes and subsequently filtered through a 30 μm cell strainer. The filtered cell suspension was centrifuged (500g, 5 minutes, 4°C) to pellet the nuclei. Nuclei were washed and filtered twice through a 40 μm cell strainer using 2% BSA in sterile PBS with 0.2 U/μl RNase inhibitor (nuclei wash). After resuspension in 500 μl nuclei wash and 900 μl 1.8M sucrose, nuclei were carefully layered on top of 500 μl of 1.8M sucrose and centrifuged (13000g, 45 min, 4°C) to remove myelin debris. The pellet was resuspended in nuclei wash and filtered again through a 40 μm Cell Strainer.

### Tissue processing and immunohistochemistry

Animals were perfused transcardially using Ringer solution (0.9% NaCl). Brain tissue was removed and immediately transferred to liquid nitrogen for subsequent RNA sequencing. For immunohistochemistry, animals where perfused using Ringer solution followed by 4% PFA solution. Brain tissue was removed and incubated for 6h in 4% PFA. Brains were cut into 40 μm coronal sections using a Thermo Scientific HM 450 microtome. Brain sections were washed with 0.1M phosphate-buffered saline (PBS) and incubated with 500 μl blocking buffer consisting of donkey serum (5 %) diluted in 1 x PBS + 0.1% Triton® X-100 for 1h at RT. Sections were then incubated with primary antibodies on a Unimax 1010 shaker at approximately 90 rmp overnight at 4°C (**Table 1**). The following day, sections were washed and incubated with corresponding secondary antibodies (**Table 2**) for 2h at RT. Sections were additionally incubated with DAPI (Sigma, 1:2000 diluted in 0.1M PBS). Sections were then mounted in 0.1M PBS onto Superfrost PlusTM microscope slides and coverslipped using Mowiol® mounting solution.

**Table 1:** Primary Antibody List.

| Antigen | Target | Host | Dilution | Company |
| --- | --- | --- | --- | --- |
| CD 13 | Pericytes | goat | 1:200 | R&D Systems |
| CD 31 | Vascular endothelial cells | rat | 1:50 | BD Biosciences |
| Fibrinogen | Blood vessel leakage | rabbit | 1:100 | abcam |
| HuNu | Human nuclei | mouse | 1:200 | EMD Millipore |
| Ki-67 | Marker for cell cycling and proliferation; tumor marker | rabbit | 1:200 | Invitrogen |
| hNanog | Pluripotency marker | goat | 1:30 | R&D Systems |
| GFAP | Astrocytes | guinea pig | 1:200 | R&D Systems |
| MAP2 | Neurons | mouse | 1:200 | R&D Systems |
| **Pax6** | Neural progenitor cells | rabbit | 1:40 | R&D Systems |
| **Iba1** | Microglia | goat | 1:200 | R&D Systems |
| **Casp-3** | Apoptotic cells | rabbit | 1:200 | Cell Signaling |
| **NeuN** | Neurons | mouse | 1:200 | Abcam |
| **hNCAM** | Graft-derived neurites | rabbit | 1:10000 | Abcam |
| **Olig-2** | Oligodendrocyte precursor cells | rabbit | 1:50 | Abcam |
| **NF-L** | Neurofilament light chain | guinea pig | 1:250 | Synaptic Systems |
| **NF-H** | Neurofilament heavy chain | rabbit | 1:100 | Sigma |
| **VIP** | Interneurons | rabbit | 1:300 | Invitrogen |
| **SST** | Interneurons | rat | 1:500 | Milipore |
| **PVALB** | Interneurons | rabbit | 1:250 | Invitrogen |
| **Synaptophysin** | Presynaptic vesicles | rabbit | 1:200 | Invitrogen |

**Table 2:** Secondary Antibody List.

| **Reactivity** | **Host** | **Conjugate** | **Dilution** | **Company** |
| --- | --- | --- | --- | --- |
| **anti-rabbit** | Donkey | Cy5 | 1:100 | *Jackson* |
| **anti-rat** | Donkey | Cy3 | 1:500 | *Jackson* |
| **anti-goat** | Donkey | AlexaFluor 488 | 1:500 | *Jackson* |
| **anti-mouse** | Donkey | Cy3 | 1:500 | *Jackson* |
| **anti-rabbit** | Donkey | AlexaFluor 488 | 1:500 | *Jackson* |
| **Anti-mouse** | Donkey | AlexaFluor 488 | 1:500 | *Jackson* |
| **Anti-goat** | Donkey | Cy3 | 1:500 | *Jackson* |
| **Anti-rabbit** | Donkey | AF647 | 1:100 | *Jackson* |
| **Anti-rabbit** | Goat | AF647 | 1:500 | *Vector Labs* |

To detect proliferation of grafted cells, 5-ethynyl-2′-deoxyuridine (EdU) labeling was performed. EdU was dissolved in sterile PBS (5 mg/ml) and injected into mice (50 mg/kg body weight) 7 days after transplantation. Brain sections were incubated for 30 min with a reaction mix buffer (1 x per well equates to 325 μl ddH_2_O, 25 μl 2M Tris, 100 μl 10mM CuSo4, 0.1 μl 10mM AlexaFluor 647, 100 μl 500 mM Ascorbic acid). Sections were washed with 0.1M PBS, incubated with DAPI, mounted onto Superfrost Plus^TM^ microscope slides, and coverslipped using Mowiol^®^ solution.

To visualize neurites extending from the transplanted cells, we performed 3,3′-Diaminobenzidine Tetrahydrochloride (DAB) staining for a human-specific neural cell adhesion molecule (hNCAM). Briefly, sections were treated with an antigen retrieval buffer at 70° for 30 minutes, followed by incubation in quenching solution for another 30 minutes. Sections were blocked (BSA (5%), Glycin (1.5%), Triton X-100 (0.25%) and Casein (1%)) and subsequently incubated with NCAM (1:10000, Abcam, #ab75813) at 4° overnight. The next day, sections were incubated with the corresponding secondary antibody (1:500, Vector Labs #BA-1000) for 2 hours at room temperature. Sections were then treated with the Avidin-Biotin-Complex (ABC) kit solution (Vector Laboratories, #PK-6100) according to the manufacturer’s protocol. The DAB staining was performed at a concentration of 0.5mg/ml DAB and 30% H2O2. Coronal sections were imaged on the Zeiss Axio Scan.Z1 slide scanner and later processed using Fiji.

### Immunostaining and analysis of whole mount muscle preparations

For immunostaining of NMJs, muscles were processed as previously described^111,112^. Briefly, EDL and the SOL muscles were carefully dissected, pinned on a Sylgard dish and rinsed with PBS. Muscles were then fixed in 4% paraformaldehyde (PFA) for 20 minutes at room temperature and subsequently washed in PBS. Muscles were cut into bundles, permeabilized in 1% Triton-X100 and remaining PFA was neutralized and blocked in 100 mM glycine in PBS, 3% BSA, 0.2% Triton-X100. Bundles were then incubated with primary antibodies against Neurofilament 200 (Ab0039, Sigma N4142, 1:6000) and Synaptophysin (Ab0736, Invitrogen MA5-14532, 1:200) followed by incubation with the corresponding secondary antibodies (Cy3, Jackson, 1:500 and Alexa488, Invitrogen, 2 μg/m). Bundles were then washed four times for 10 min in PBS and mounted with ProLong™ Gold antifade (Invitrogen). Images were taken on either a wide field IX83 microscope (Olympus) or a confocal LSM 700 (Zeiss).

To analyse NMJ innervation status as well as NMJ area, z-stacks were projected in 3D using the ImageJ software. Only NMJs with clearly visible presynaptic axons and terminals were scored. At least 20 NMJs were scored per mouse and per muscle for innervation status and at least 12 for NMJ area measurements. NMJs were categorized into the following groups: poly-innervated (when at least two sources of innervation, either axons or thin sprouts, contacted one NMJ), partially innervated (when one or two presynaptic inputs contacted one NMJ) and denervated (with no presynaptic input at all). To measure the NMJ area, the perimeter circumscribing the postsynaptic staining was drawn by hand and the enclosed surface area was measured using ImageJ.

### Axonal tracing

At 28 days post-transplantation, 1 week prior to perfusion, a subset of animals (NPC: *n*=6, vehicle, *n*=4) were injected with 0.2 µl of the anterograde axonal tracer biotinylated dextran amine (BDA, molecular weight 10 000, 0.1 µg/µl; Molecular Probes) at 0.2 µl/min into the ipsilesional stroked cortex at: AP: + 0.5 / MV: + 1.5 / DV: - 0.6 (mm relative to bregma). The needle was left in place for 5 minutes, and then retracted slowly. At 35 days post-transplantation, animals were perfused, and brains were processed for immunohistochemistry. BDA was detected using streptavidin-conjugated antibody (Alexa Fluor 647, 1:200, Thermo Fisher Scientific). BDA-positive fibers were determined from regions adjacent to the stroke core in two brain slices per animal at pre-defined landmarks. ImageJ software was used to determine the are fraction of BDA-positive area above a set threshold; the values were then normalized to the number of BDA-positive cell bodies.

### Microscopy stroke area/volume quantification

Imaging of brain sections was performed with a *Leica SP8 laser confocal microscope* with 10x and 63x objectives, and images were processed using *Fiji* (ImageJ) as previously described^108^. For stroke and graft size analysis, 40 μm coronal sections stained with HuNu (graft) and GFAP (lesion size) were imaged on the Zeiss *Axio Scan.Z1* slide scanner and later processed using *Fiji*. The lesion area of each coronal section was measured by manually drawing a polygonal area around the stroke region as closely as possible. Brain sections were identified according to stereotaxic coordinates (distance relative to Bregma) using a mouse brain atlas^113^. A custom-made RScript was used to re-scale the image and enhance contrast. Additional parameters (height, width) were recorded for graft size analysis. To assess lesion- and graft size volume, length, width, and area were measured for each brain section and ‘stacked together’ to create a 3-D model. Lesion- and Graft volume was approximated as an elliptical oblique cone:

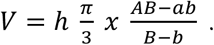

### Histological quantification of vasculature, microglia and astrocytes

All analysis steps were conducted in a peri-infarct region, referred to as ischemic border zone (IBZ) adjacent to the stroke core that extended up to 300 μm. Post-ischemic angiogenesis was assessed using the software FIJI (ImageJ, version 2.1.0/1.53c) and previously established script that allows to automatically calculate (1) vascular density, (2) number of branches and junctions and (3) the length of blood vessels^79^. The intensity of the glial scar and inflammation signal was determined by analyzing three brain sections per animal, which were immunostained for GFAP and Iba1. Images were converted to 8-bit format and subjected to thresholding using mean grey values obtained from regions of interest (ROIs) in the unaffected contralateral cortex to create a binary image. The cumulative area of reactive gliosis or inflammation in and surrounding the ischemic lesion was then computed.

Morphology of Iba1^+^ cells was performed using the skeletal analysis plugin and/or the Sholl Analysis plugin in Fiji (ImageJ). In brief, 63x z-stack images were projected using a maximum intensity projection. The brightness/contrast of the z-stack was then adjusted to best visualize the branches of the Iba^+^ microglia. Images were set to *Grayscale* and subsequently converted to 8-bit format and subjected to thresholding using mean grey values obtained from regions of interest (ROIs) in the unaffected contralateral cortex to create a binary mask. Circularity was calculated using the built-in plugin *Analyze Particles* (ImageJ). A circularity value of 1.0 indicates a perfect circle, while values approaching 0.0 indicate increasingly elongated polygons.

For calculating the branching and the ramification index, images were skeletonized using the Analyze Skeleton plugin^114^ followed by the Sholl analysis plugin^115^. A radius was drawn from the center of the cell body to the end of the longest branch to set the upper and lower limit for concentric circle placement. The first circle was set as close to the edge of the cell body as possible to ensure the cell body was not counted as an intercept on the circle. The distance between each circle was set at 2 µm for all cells.

### Cell counting

Cell counting was completed with Fiji (ImageJ). 63x images were imported and Z-stacked using a ‘standard deviation’ projection. The ‘cell counter’ plugin was used to count cells manually. Brain sections were double stained with HuNu antibodies or analyzed with endogenous GFP, in addition to the staining (Nanog, Pax6, MAP2, GFAP, Ki67, EdU) to be quantified. Cells were counted separately and independent of each other for each section. DAPI was co-stained and used as a control measure to ensure that cells were not mistaken for cell debris. The total of all characterized cells (Nanog, Pax6, MAP2 and GFAP) was set to 100%.

### Rotarod performance test

Mice were first acclimatized to the rotarod in a habituation phase 4 days before stroke induction by placing them on a rotating rod at 4 rpm for 60 seconds. The test phase consisted of three trials separated by approximately 15 min inter-trial intervals, which acted as a recovery phase. The apparatus was set to ‘accelerating mode’ with a starting velocity of 5 rpm to a maximal speed of 50 rpm in 300s. Latency (=amount of time that lapses between begin and end of behavioral recording) recorded the time point at which each mouse fell off the rod and onto an underlying panel that stopped the counter. In addition, latency was also recorded if a mouse completed two full passive rotations in a row.

### Horizontal ladder walk

The horizontal ladder walk identifies stroke-related gait abnormalities across a specific time period. Animals were placed on a modified MotoRater 303030 series with a horizontal ladder rung made of metal (length: 113 cm, width: 7 cm) and were trained to cross from one end of the ladder to the other. The structure was made of a clear Plexiglas basin (156 cm long, 11.5 cm wide, 11.5 cm high) and was equipped with two perpendicularly arranged mirrors to allow simultaneous image recording from different angles. To prevent habituation to a specific bar distance, bars were irregularly spaced (1-4 cm). For the behavioral test, a total of three runs per animal were recorded. A misstep was defined as the mouse’s toe tips reaching 0.5 cm below the ladder rung. The error rate was calculated by errors / total steps* 100. The movement pattern was recorded by a high-definition video camera (GoPro Hero 7) at a resolution of 4000 x 3000 and a rate of 60 frames per second. Videos were cut and formatted with Adobe Premier and analyzed with DeepLabCut and RStudio.

### Horizontal runway test

A runway test was performed to assess whole-body coordination during locomotion. As in the horizontal ladder walk, the modified MotoRater 303030 series was used, with exclusion of the ladder. Mice were recorded crossing the runway with a high-definition video camera (GoPro Hero 7) at a resolution of 4000 x 3000 and a rate of 60 frames per second. In a habituation phase, each animal was trained in two daily sessions until they crossed the runway voluntarily (without external intervention) at a constant speed. For behavioral test recording, each animal was individually placed on one end of the basin and was allowed to walk for 3 minutes.

### DeepLabCut

Video recordings were processed by DeepLabCut^TM^ (DLC, v. 2.1.5), an open-source software that enables the computation of 2D and 3D marker-less pose estimates to track limb movements of animals. A small subset of camera frames served as a training dataset and were manually labeled as accurately as possible for each test, which was used to train an artificial network. Once the network was trained adequately, recordings of behavioral experiments were fed into DLC for automated tracking. The artificial network algorithm could then predict the marker locations in new recordings. 2D points can be further processed for performance evaluation and 3D reconstruction. Data was processed with R. Video pixel coordinates for the labels produced by DLC were imported into R Studio (Version 4.04 (2021-02-15) and processed with custom scrips that can be assessed here: https://github.com/rustlab1/DLC-Gait-Analysis.

### Single-nucleus RNA sequencing

Single-nucleus RNA sequencing (snRNAseq) was performed as previously described.^37^ For droplet-based library preparation, isolated nuclei obtained from stroked and non-stroked mouse cortices were loaded onto the Chromium platform from 10x Genomics according to manufacturer’s instructions (10X Genomics). RNA molecules were captured for amplification using the Chromium 3 Reagent Kits v3. The libraries were sequenced using an Illumina sequencer. Sample demultiplexing, barcode processing and single-cell counting was performed using the Cell Ranger Single-Cell Software Suite (10x Genomics).

### Clustering and annotation of mouse and human cell types

Alignment and gene quantification of single-nucleus RNA sequencing (snRNAseq) data were carried out using Cellranger v3.1.0, adhering to the default settings, and using the human and mouse (GRCh38 & mm10) 2020-A datasets as reference. Cells exhibiting more than 5% mitochondrial gene expression and less than 500 nFeature_RNA were excluded. Gene counts underwent normalization and scaling to account for the total unique molecular identifier counts per barcode using Seurat v5.0.1^116^. All analysis steps were performed according to the guidelines in the Seurat package^116^. For cell clustering, the initial 30 principal components derived from principal component analysis were employed to identify neighbors using the FindNeighbors function, followed by clustering using the FindClusters function. Dimensionality reduction via Uniform Manifold Approximation and Projection (UMAP) was performed using the RunUMAP function. Classification of distinct cell types was informed by established cell-type markers from human and mouse reference datasets^38–40^. The datasets were categorized into Glut (glutamatergic cells), GABA (GABAergic cells), Astro (astrocytes), Oligo (oligodendrocytes), and NN (non-neural cells). The identification of cell type–specific marker genes was performed using the FindMarkers function, with all parameters set to default. Genes with Bonferroni correction adjusted p-value <0.05 were considered marker genes.

### Inference of cell-cell communication networks

Differential cell–cell interaction networks were reconstructed using CellChat version 2.1.0^46,47^ for both human and mouse datasets. In brief, DifferentialConnectome was applied to the Seurat objects (version 5.01), which contained integrated data of human and mouse datasets. Mouse data was transformed to human orthologues prior analysis. The total numbers of interactions and interaction strengths were calculated using the compareInteractions function. Network centrality scored were calculated using the netAnalysis_computeCentrality function. All analysis were performed according to the tutorial called ‘Full tutorial for CellChat analysis of a single dataset with detailed explanation of each function’ on the GitHub page. For pathway analysis, signaling pathways were grouped based on functional and structural similarity as defined in CellChat. Functional similarity in CellChat refers to pathways that regulate related biological processes, whereas structural similarity is based on shared ligand-receptor families or overlapping downstream signalling components. The terms *Sender*, *Receiver*, *Mediator*, and *Influencer* represent distinct roles in intercellular communication inferred from CellChat’s network analysis: **Sender (Source) Cells**: Cells that secrete ligands, initiating the signaling process. **Receiver (Target) Cells**: Cells that express the corresponding receptors and respond to the signals. **Mediator Cells**: Cells that facilitate or amplify communication between sender and receiver cells, acting as intermediaries in the network. **Influencer Cells**: Cells that modulate the strength or efficacy of a signaling pathway, shaping how communication occurs even if they are not direct senders or receivers.

### Statistical analysis

Statistical analysis was performed using RStudio (Version 4.04). Sample sizes were designed with adequate power in line with previous studies from our group and relevant literature. Generated data was tested for normal distribution using the Shapiro-Wilk test. Normally distributed data was tested for differences with an unpaired two-sided t-test to compare changes between two treatment groups (i.e differences between ipsi- and contralesional sides). Non-normally distributed data was transformed using natural logarithms or exponential transformation before completing statistical tests to assume normal distribution. The significance of mean differences between normally distributed multiple comparisons (i.e. longitudinal data of treatment groups) was assessed using ANOVA with post-hoc analysis (estimated marginal means). Data is expressed as mean ±SD; statistical significance was defined as *p < 0.05, **p < 0.01, and ***p < 0.001.

### Data availability

All summarized data and statistical analysis are accessible in the supplementary materials (**Suppl. Table 1**). Single nucleus RNAseq data are accessible via NCBI GEO (GSE259430). Single nucleus RNAseq datasets can be explored via Shinyapp: https://rustlab.shinyapps.io/Cell-Therapy-Stroke-Atlas/ and https://rustlab.shinyapps.io/cell-graft-stroke/.

## Acknowledgement

This work is supported by the funding from Swiss 3R Competence Center (OC-2020-002) and the Swiss National Science Foundation (CRSK-3_195902) and (PZ00P3_216225) to RR and from the Neuroscience Center Zurich to CT. In addition, the authors acknowledge support from the Mäxi Foundation.

## Author contribution

RZW, CT, RR contributed to overall project design. RZW, KK, RR contributed to the design of snRNAseq experiments. RZW, BAB, NHR, PP, RR conducted and analyzed in vivo experiments. CB, KJZ, DU, DM, SP conducted and analyzed in vitro experiments. RZW, AB, MZ, RR performed nuclei isolation and snRNAseq experiments. RR analyzed snRNAseq experiments. RZW, CT and RR made figures. MG provided an iPS cell line. SL and MAR performed neuromuscular junction analysis. RMN, CT, RR supervised the study. RZW, KK, CT, RR wrote and edited the manuscript with input from all authors. All authors read and approved the final manuscript.

