## Supplemental Data for "Neural xenografts contribute to long-term recovery in stroke via molecular graft-host crosstalk"

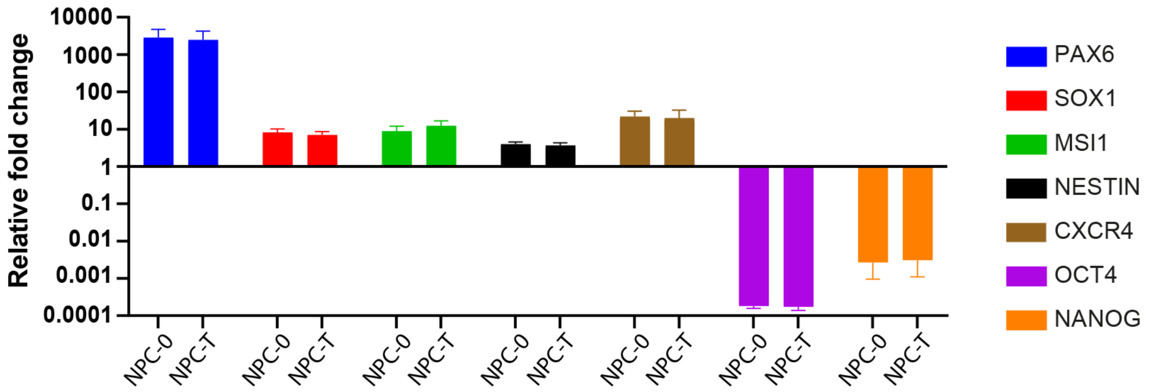


**NPC-0:** NPCs untransduced, passage 8

**NPC-T:** NPCs transduced with **GFP** and **luciferase** construct, passage 8

**Suppl. Fig. 1: Characterization of NPCs.** Gene expression of NPC marker (PAX6, SOX1, CXCR4, MSI1, NESTIN, CXCr4) and pluripotency marker (NANOG and OCT4) in untransduced NPCs (NPC-0) and NPCs transduced with dual-reporter (NPC-T), measured by qPCR, presented as fold change over iPSCs.

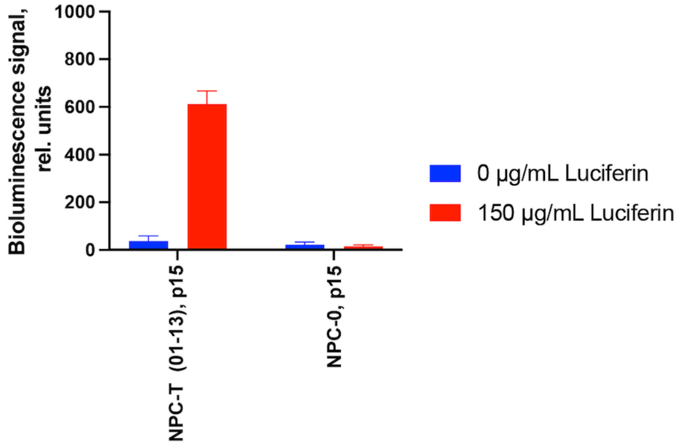


**NPC-0:** NPC untransduced, passage 15

**NPC-T:** NPC (01-13) transduced with **GFP** and **luciferase,** passage 15

**Suppl. Fig. 2: In vitro validation of the luciferase reporter system.** Luminescence assay showing the luminescence signal in NPCs, transduced (left) and untransduced (right).

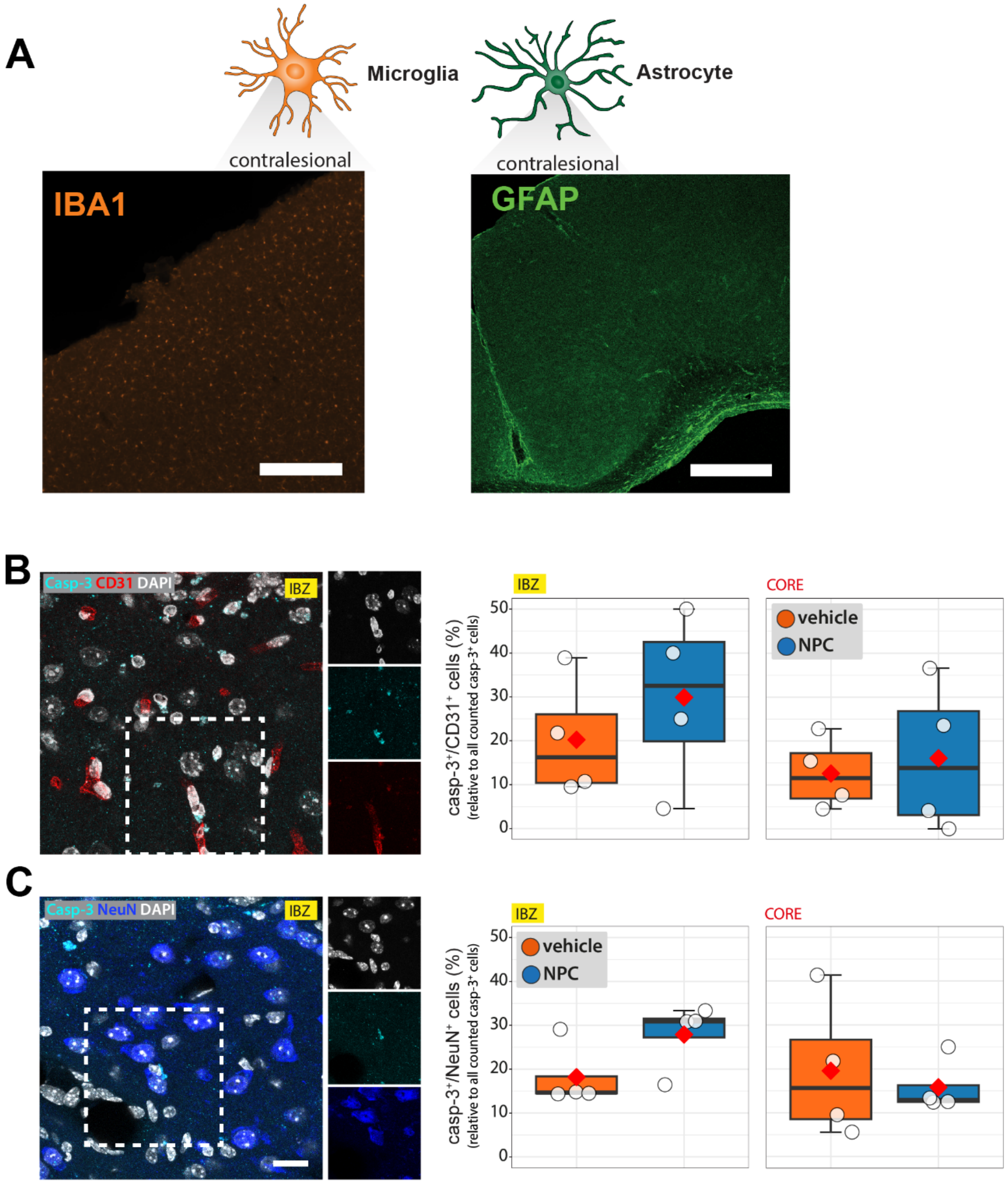


**Suppl. Fig. 3: Histological analysis of microglia, astrocyte expression in the contralesional hemisphere and characterization of Casp-3$^{+}$ cells in the IBZ and the core region**. (A) Representative images of contralesional microglia (**Iba1**, orange) and astrocytes (**GFAP**, green) staining. Scale bar: 100 µm. (B) Immunofluorescent staining of a coronal brain section with Casp-3, CD31 and DAPI and (right) quantification of Casp-3/CD31+ cell count relative to all counted Casp-3+ cells in the IBZ and the core region. (C) Immunofluorescent staining of a coronal brain section with Casp-3, NeuN and DAPI and (right) quantification of Casp-3/NeuN+ cell count relative to all counted Casp-3+ cells in the IBZ and the core region. Scale bar = 10µm. Data are shown as mean distributions where the red dot represents the mean. Boxplots indicate the 25% to 75% quartiles of the data. Each dot in the plot represents one animal. Significance of mean differences was assessed using an unpaired t-test (vehicle vs. NPC).

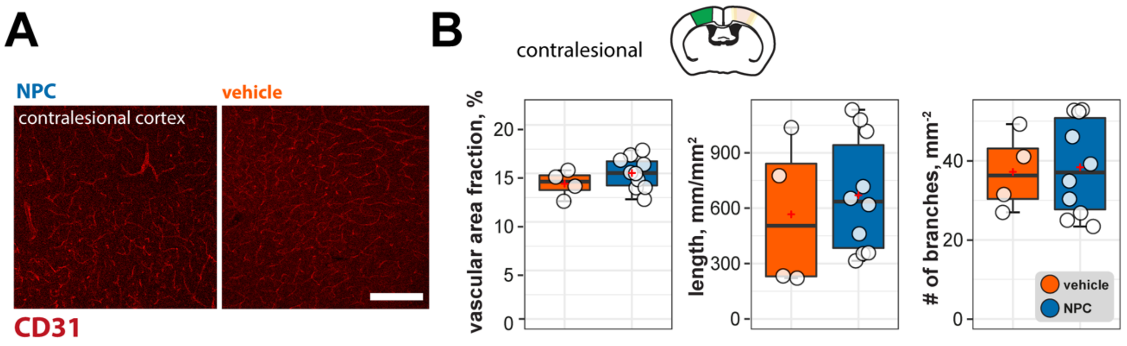


**Suppl. Fig. 4: Vascular changes in the contralesional cortex in NPC- and vehicle-receiving animals.** (A) Immunofluorescence staining of coronal brain sections with CD31 to label blood vessels on contralesional hemisphere. Scale bar: 100 µm. (B) Analysis of vasculature density (vascular area fraction, number of branches, vessel length) in the contralesional cortex. Data are shown as mean distributions where the red dot represents the mean. Boxplots indicate the 25% to 75% quartiles of the data. For boxplots: each dot in the plots represents one animal. Significance of mean differences was assessed using an unpaired t-test (vehicle vs. NPC). In B, n=10 (vehicle) and n=11 (NPC) mice per group.

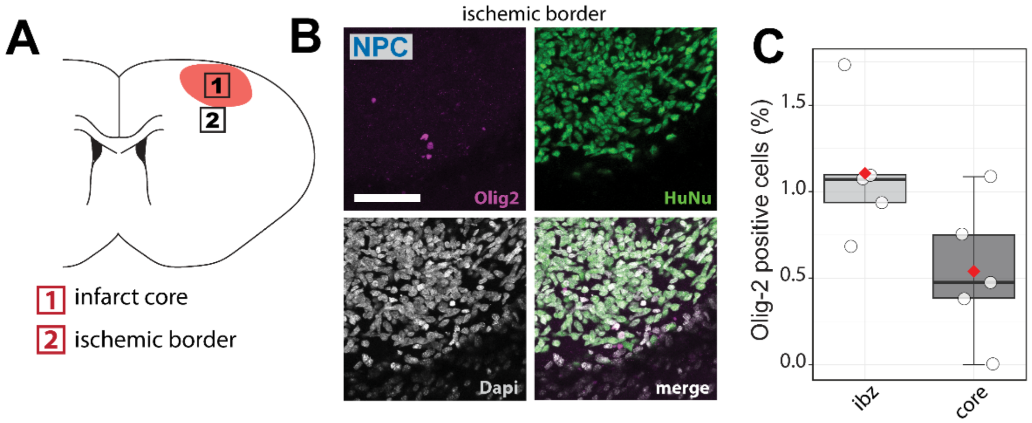


**Suppl. Fig. 5: Fate of transplanted NPCs 35 days post-transplantation.** (A) Illustration showing the two brain areas (infarct core, ischemic border zone) analysed for Olig2/Hunu double-positive cells. (B) Immunofluorescent staining of a coronal brain section with Olig2, HuNu and Dapi and (C) quantification of Olig2/HuNu$^{+}$ cell count relative to all counted HuNu$^{+}$ cells. Scale bar = 50µm. Data are shown as mean distributions where the red dot represents the mean. Boxplots indicate the 25% to 75% quartiles of the data. Each dot in the plot represents one animal. Significance of mean differences was assessed using an unpaired t-test (ibz vs. core).

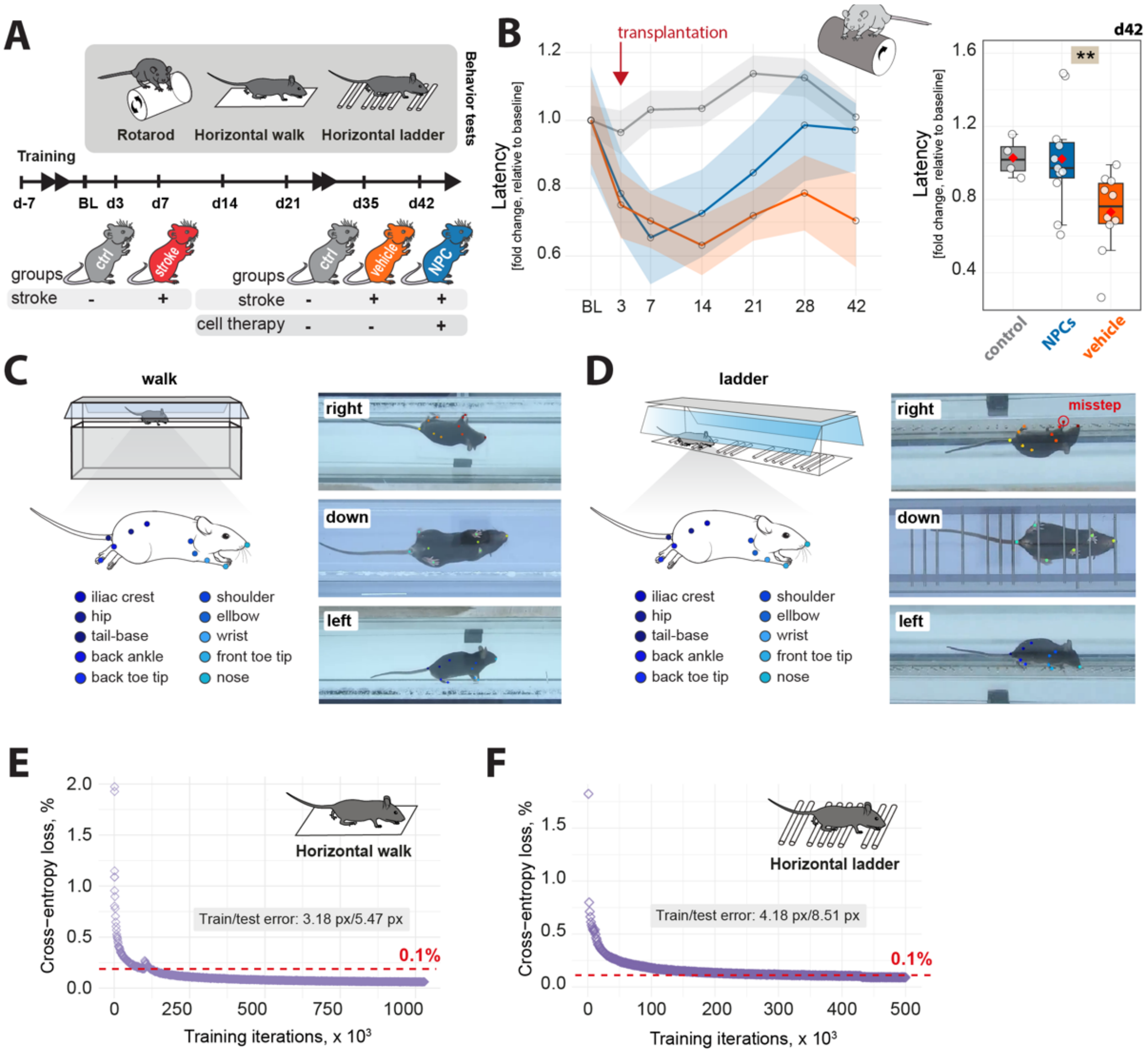


**Suppl. Fig. 6: Increased recovery of motor function for mice receiving NPCs.** (A) Schematic representation of behavioral tests of different treatment groups (non-stroked vs. stroked; non-stroke and vehicle treatment vs. stroked and vehicle treatment vs. stroked and cell therapy). (B) Analysis of rotarod performance data. Latency is defined as x-fold change in performance relative to baseline. (C) Anatomical landmarks of mice for pose estimation during gait analysis in horizontal walk and (D) during horizontal ladder rung walk from all three perspectives (right, below, left). (E) Training efficiency of neural networks for horizontal walk and (F) horizontal ladder rung. Data are shown as mean distributions with the red dots representing the mean. Line graphs are plotted as mean ± sem. For boxplots: each dot represents one animal. Significance of mean differences was assessed using an unpaired t-test (stroke vs non-stroke) or Tukey's HSD. In B n=10 (vehicle), n=11 (NPC) and n=4 (ctrl) mice per group. Asterisks indicate significance: **p < 0.01.

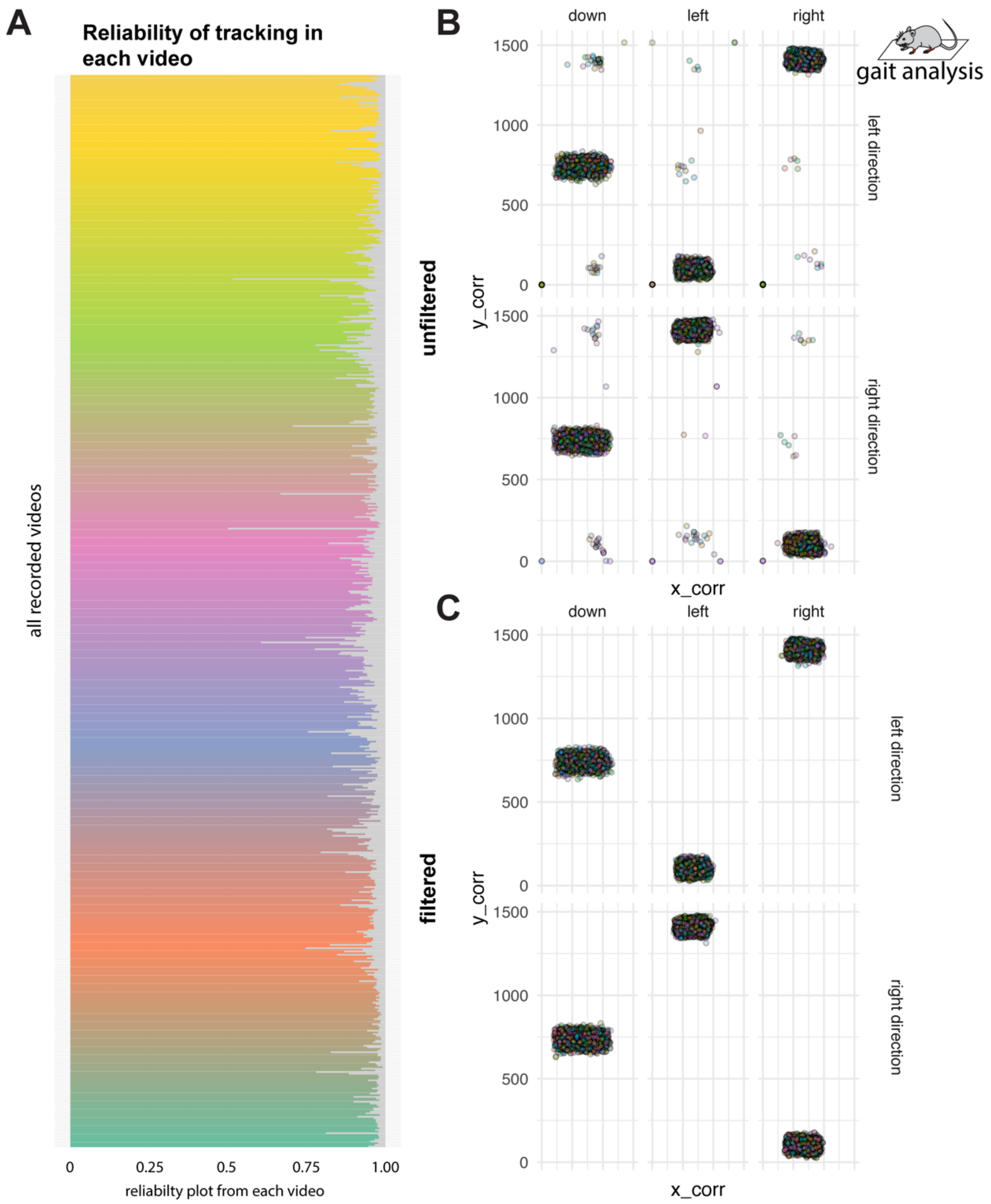


**Suppl. Fig. 7: Reliability of tracking for behavioral gait analysis.** (A) Probability of reliably tracked labels (>95% reliability) to all tracking labels for each video. (B) Dotplot of individual labels from three perspectives before and (C) after filtering for outliers.

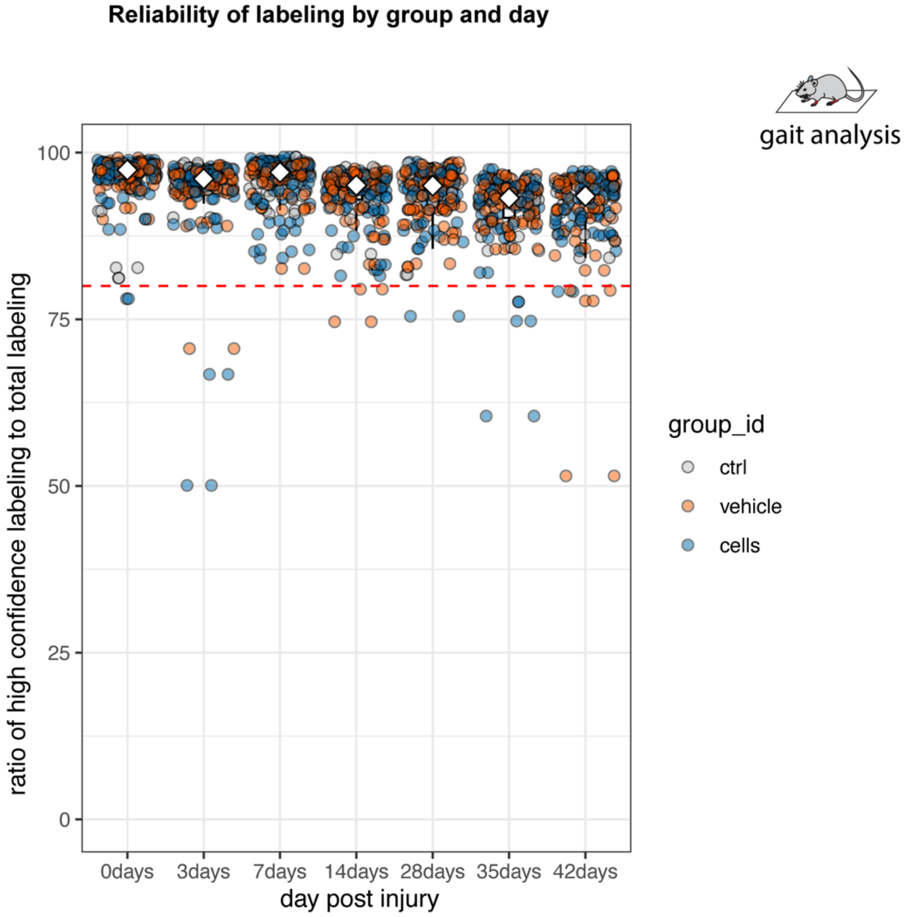


**Suppl. Fig. 8: Reliability of labeling by group and day.** Comparable ratio of reliable labels (>95% accuracy) to all labels was observed across treatment groups and days post-injury.

**Representative gait profile of three uninjured mice**

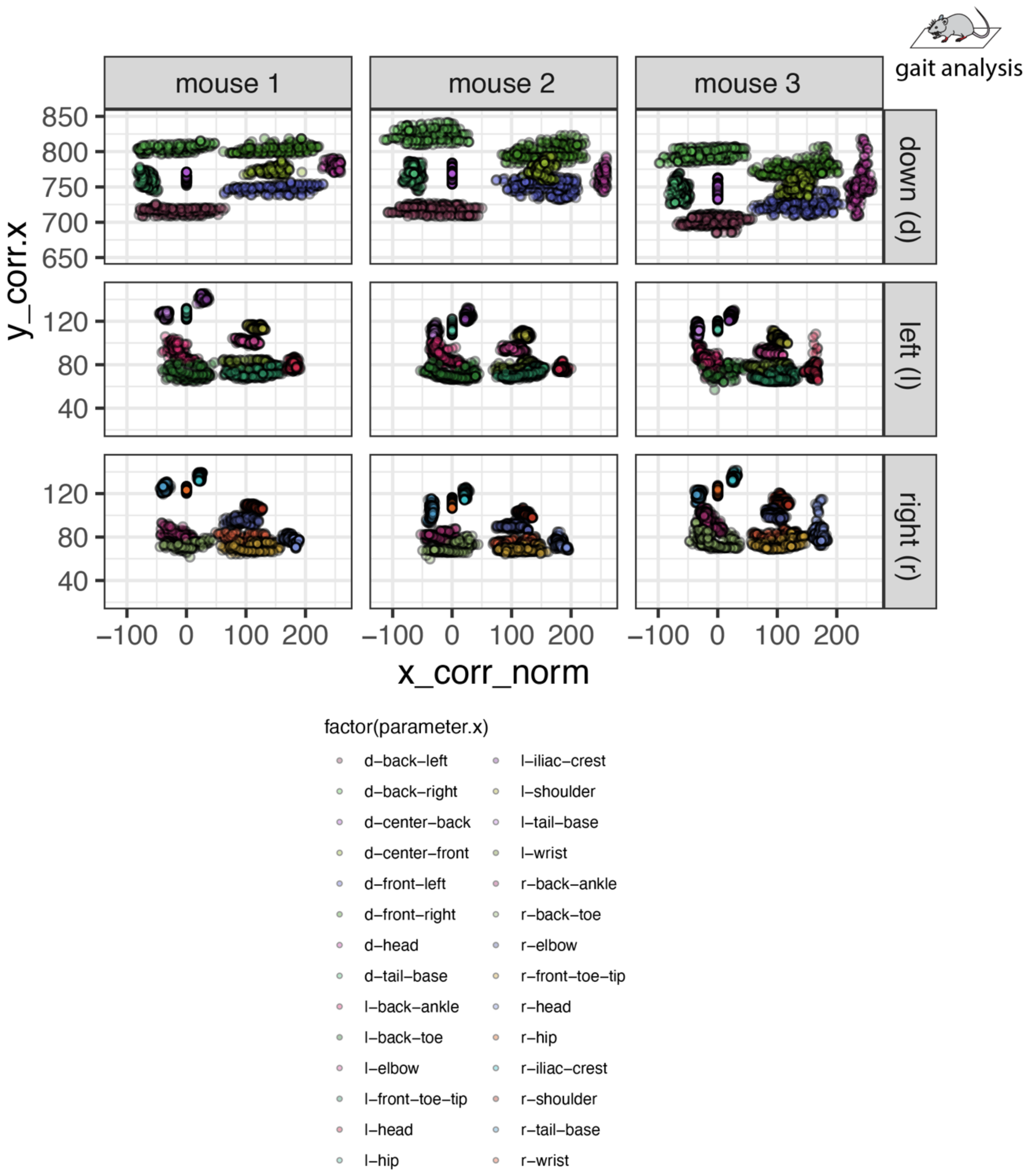


**Suppl. Fig. 9: Mouse tracking for behavioral analysis**. Representative profiles of three randomly assigned mice at baseline tracked from left, bottom and right perspective for each individual body part.

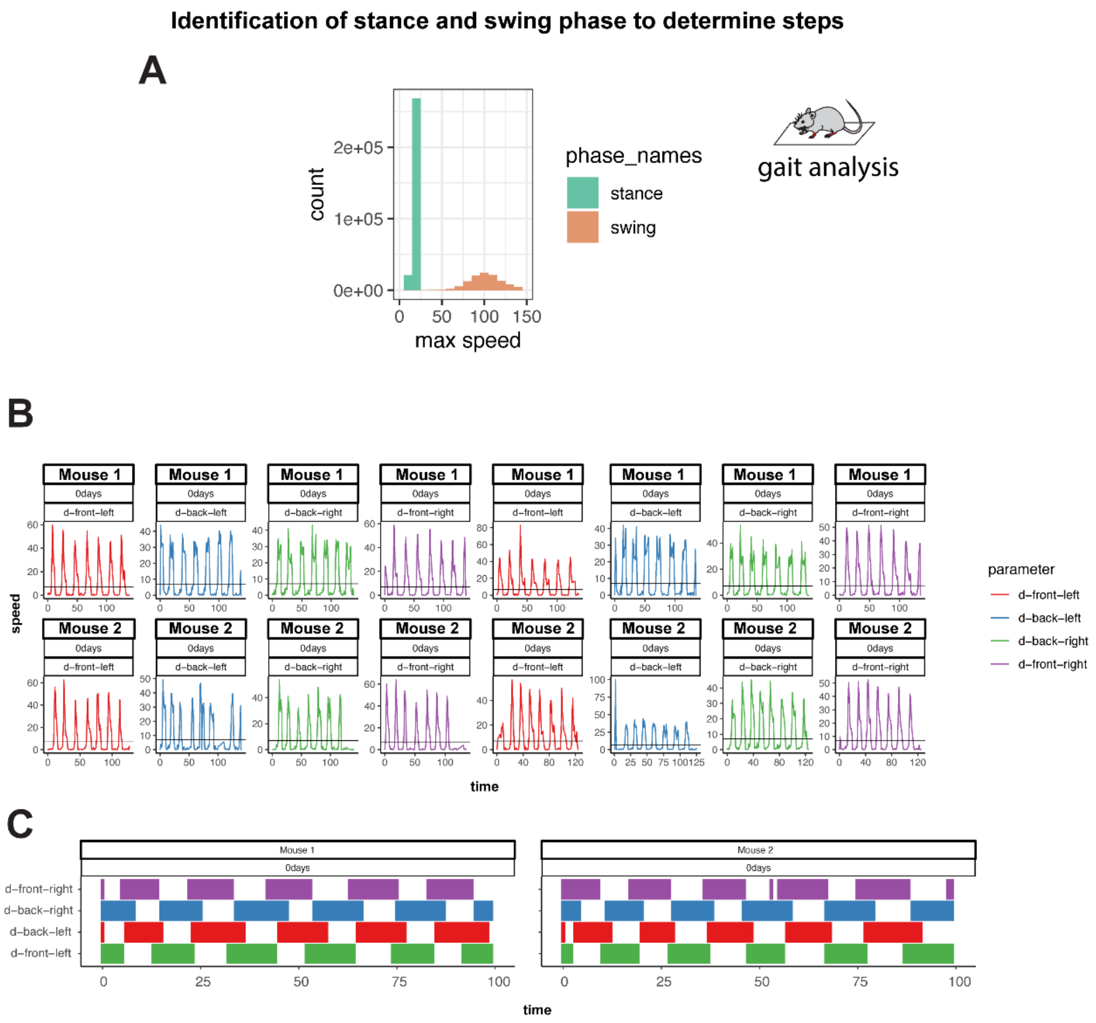


**Suppl. Fig. 10: Validation of behavioral tracking**: (A) Tracking speed of paws and separation between stance and swing phase in two randomly assigned mice at baseline. (B) Representative plots determining individual steps based on speed of paws at baseline. (C) Representative footstep profiles from two randomly assigned mice at baseline.

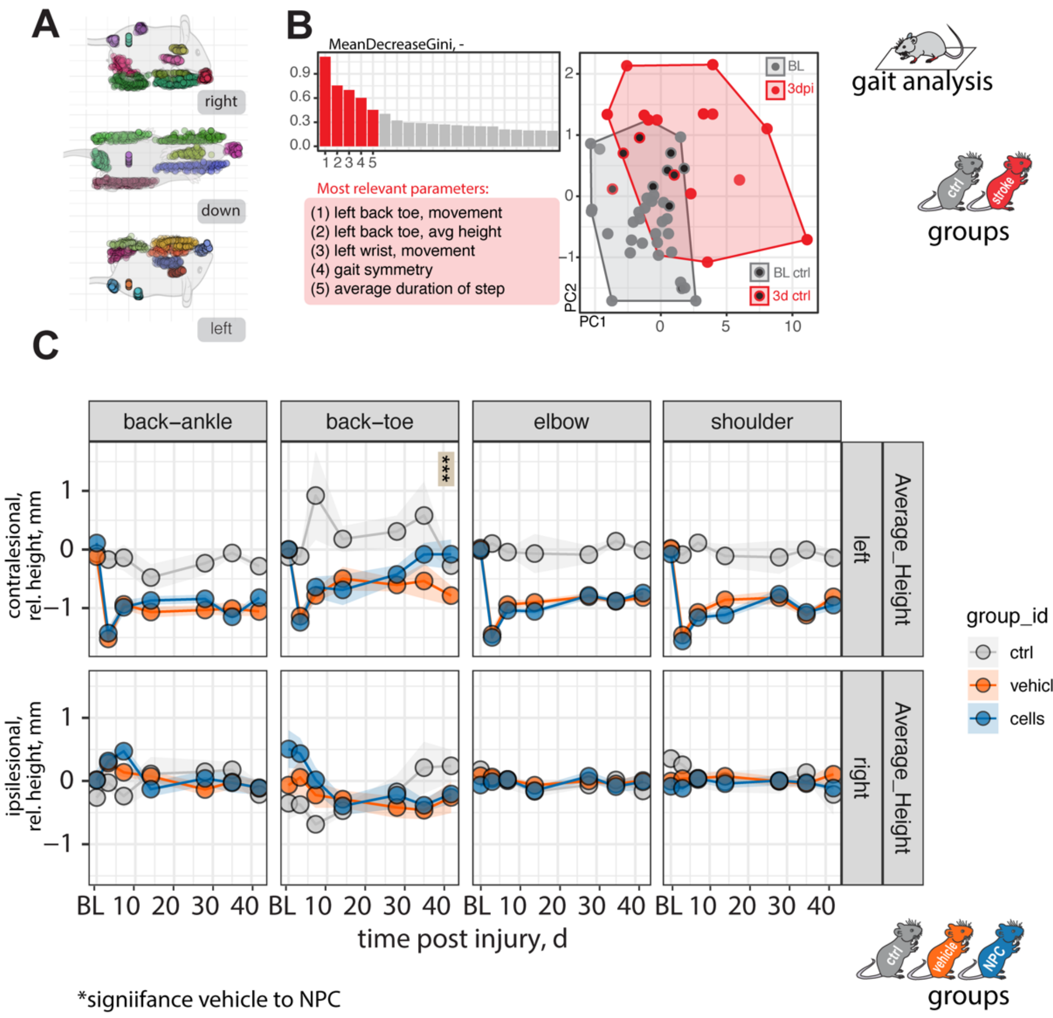


**Suppl. Fig. 11: Gait-analysis to assess behavioral recovery. (**A) Representative gait profile in a mouse (B) Left: Random Forest classification of most important parameters to separate ctrl and stroke at 3dpi. Right: Principal component analysis of most relevant parameters at 3 dpi. (C) Relative height of back ankle, back toe, elbow, and shoulder at baseline and 3, 7, 14, 21, 28 and 35 dpi. Line graphs are plotted as mean ± sem. Asterisks indicate significance: ***p < 0.001.

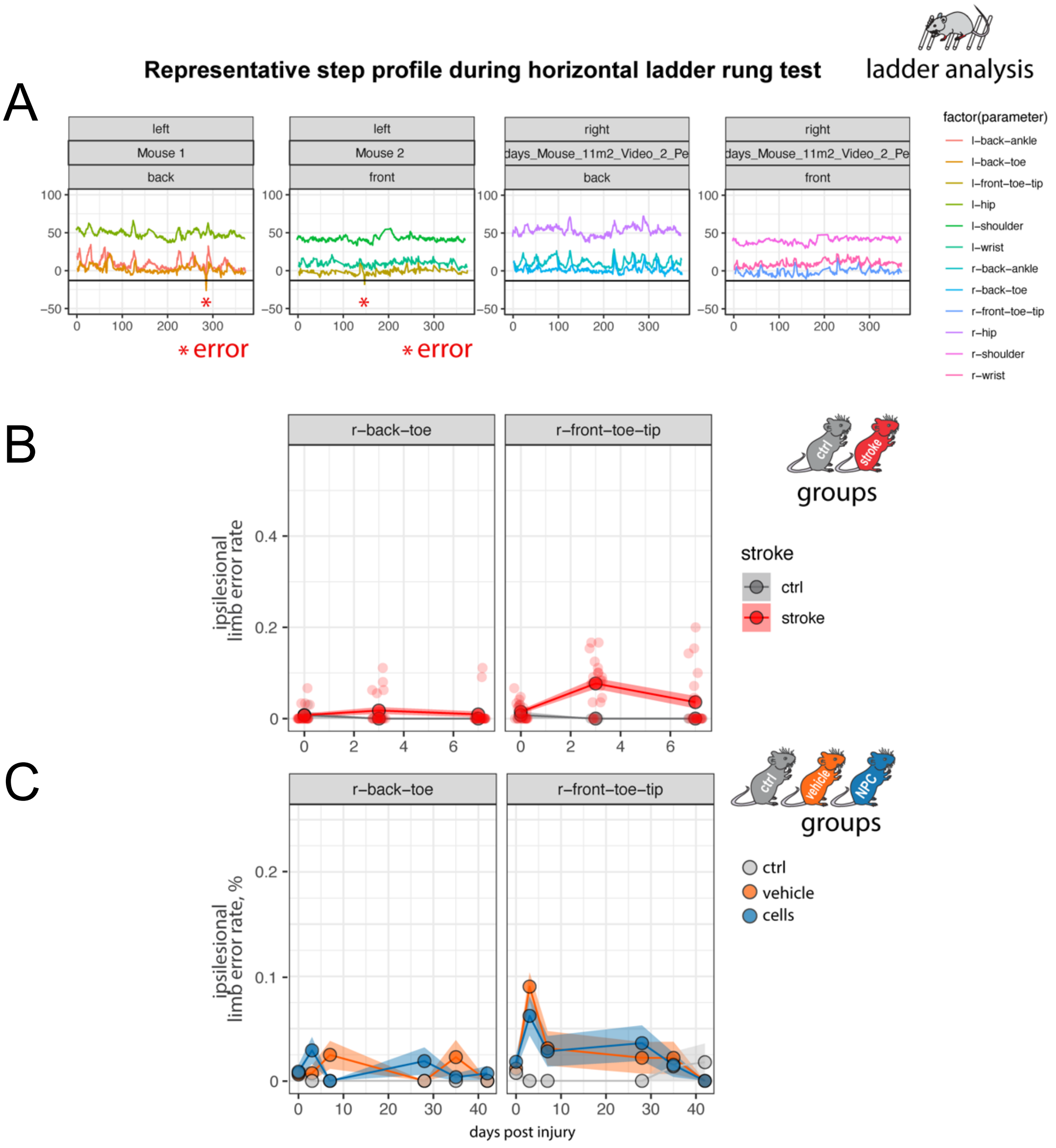


**Suppl. Fig. 12: Analysis of horizontal ladder rung test**. (A) Representative profiling of a front and back toes during the horizontal ladder rung text. (B) Overall rate of missteps to total steps in ipsilesional forelimbs and ipsilesional hindlimbs. (C) Overall rate of missteps to total steps in ipsilesional forelimbs and ipsilesional hindlimbs at baseline and 3, 7, 14, 21, 28 and 35 dpi. Missteps were detected by the back toe tips and front toe tips dipping below a defined threshold. Line graphs are plotted as mean ± sem. In B, n=21 (stroke) and n=4 (no stroke) mice per group. In C, n=10 (vehicle), n=11 (NPC) and n=4 (ctrl) mice per group.

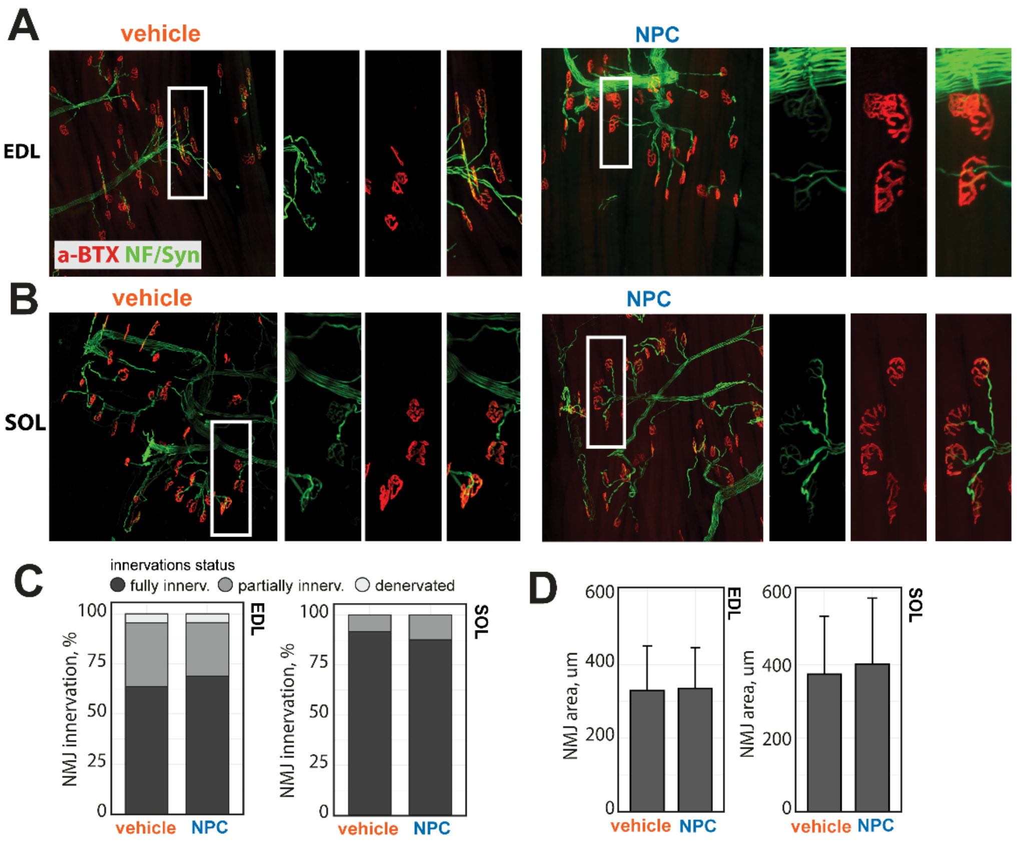


**Suppl. Fig. 13: Structural alterations in NMJs after vehicle or NPC treatment**. (A) Representative images of NMJs from EDL (A) and SOL (B) from vehicle- and NPC-treated mice. Immunofluorescence assay with α-bungarotoxin (BTX) (red, for acetylcholine receptor) and anti-synaptophysin (Syn) and anti-neurofilament (NF) antibodies (both green, for nerve terminals) used for analysis of NMJ area and innervation status. (C) Barplot showing the proportion of fully innervated, partially innervated, and denervated endplates in EDL (left) and SOL (right). (D) Quantification of NMJ area in EL and SOL muscles of vehicle- and NPC-treated animals. The area of BTX-labelled AChR represents the NMJ area. Data are plotted as mean ± sem. In C and D, we used N=3 mice per group (total N=6).

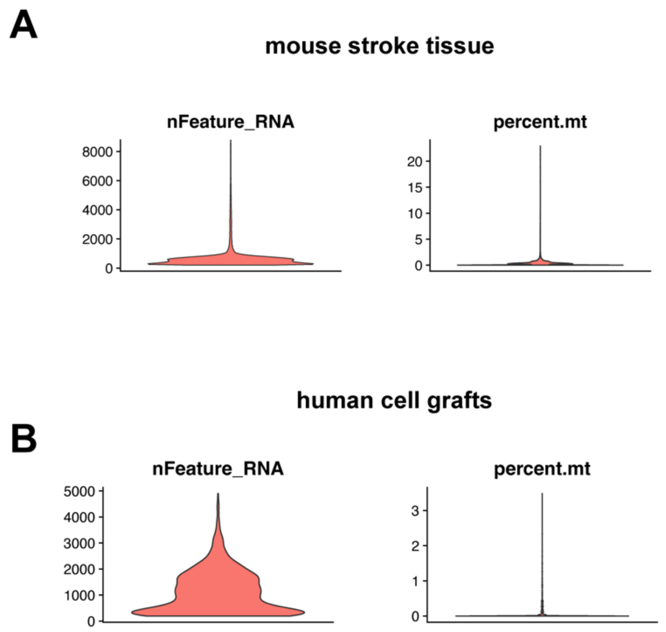


**Suppl. Fig. 14: Quality control of single nucleus RNA sequencing.** (A) Quality control of nuclei from stroke-injured mouse tissue indicating number of unique features and percentage of mitochondrial RNA. (B) Quality control of nuclei from human cell grafts indicating number of features and percentage of mitochondrial RNA.

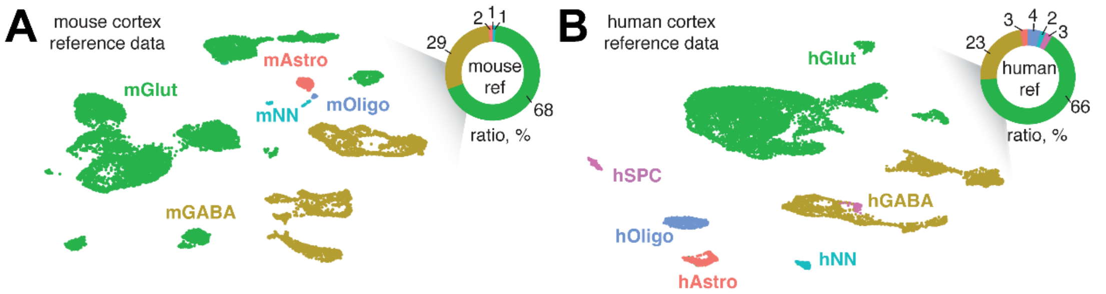


**Suppl. Fig. 15: UMAP representation of mouse and human reference datasets.** (A) Uniform Manifold Approximation and Projection (UMAP) of 1.1M cells from reference adult mouse whole cortex from Allen brain atlas colored by cell type. (B) UMAP of 76,533 nuclei from reference adult human cortex from Allen brain atlas colored by cell type.

reference mouse brain marker validation

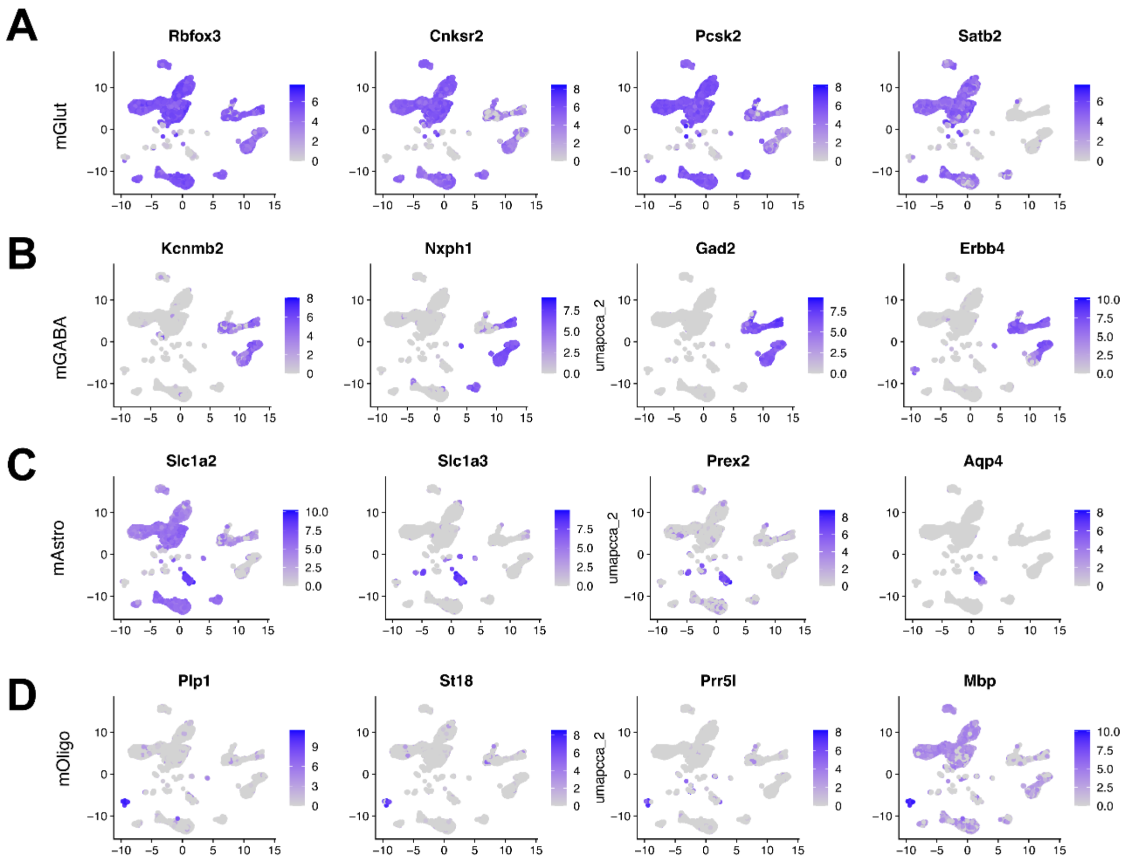


**Suppl. Fig. 16: Mouse reference dataset marker validation.** (A-D) Feature plots of gene expression for specific cell type markers in reference mouse brain dataset: glutamatergic neurons (mGlut, A), GABAergic neurons (mGABA, B), astrocytes (mAstro, C), and oligodendrocytes (mOligo, D).

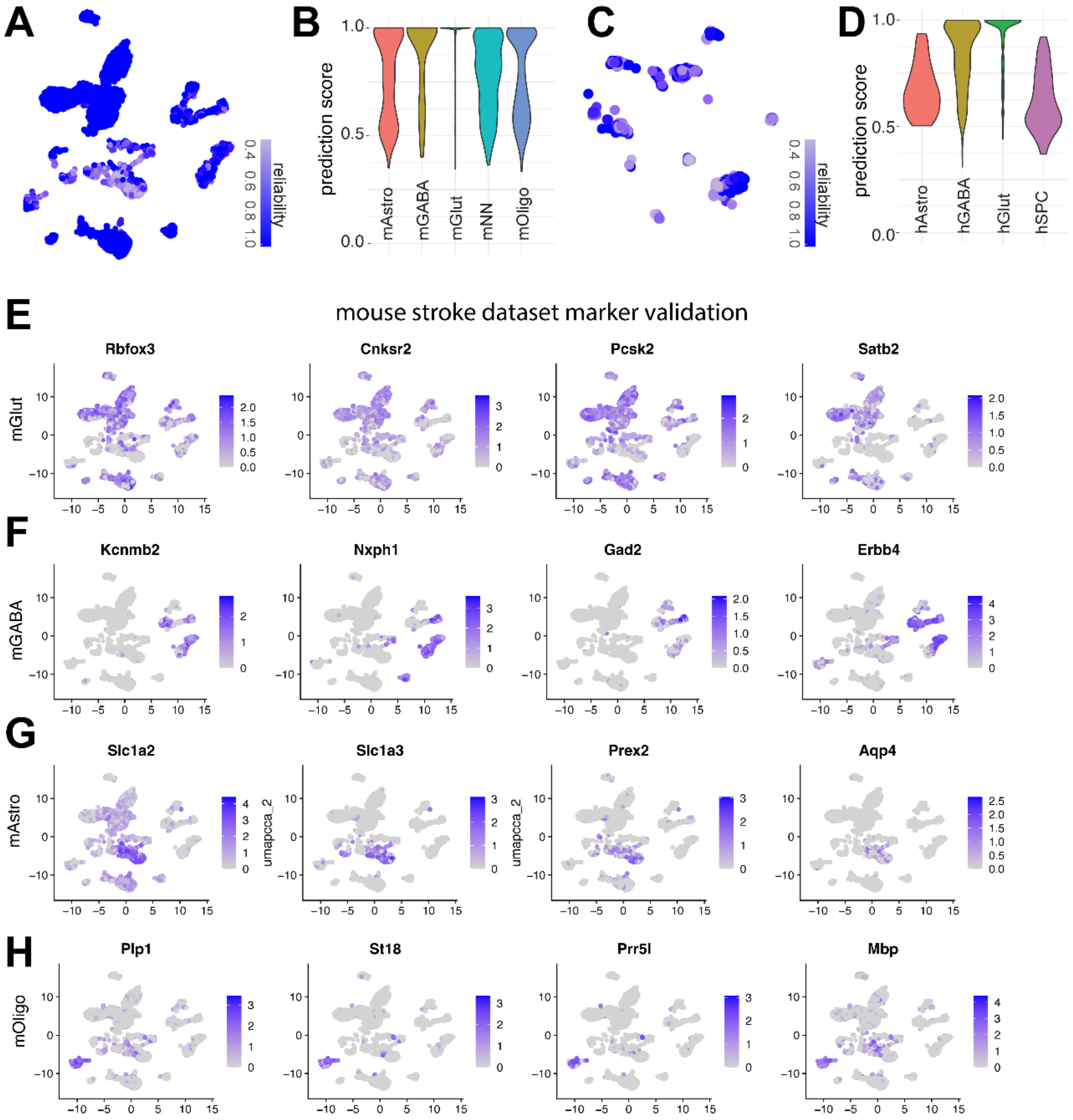


**Suppl. Fig. 17: Mouse stroke dataset marker validation.** (A, B) Reliability feature plot and prediction score of 83’058 stroke mouse tissue (n =5), (C, D) Reliability feature plot and prediction score of 1149 transplanted human cell grafts (n = 5). (E-H) Feature plots of gene expression for specific cell type markers in mouse stroke brain: glutamatergic neurons (mGlut, E), GABAergic neurons (mGABA, F), astrocytes (mAstro, G), and oligodendrocytes (mOligo, H).

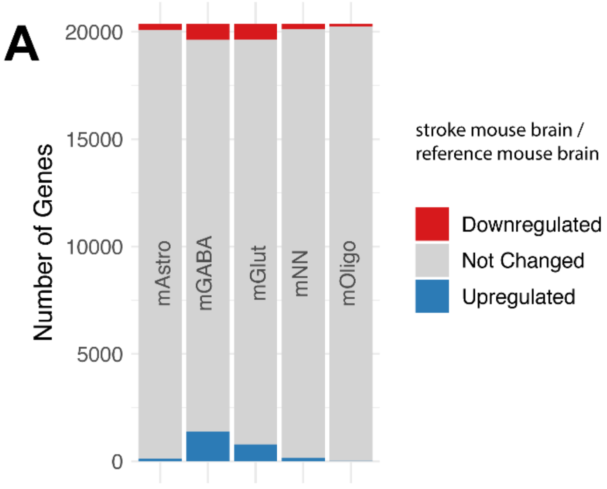


**Suppl. Fig. 18: Changes in gene expression in host brain cells after stroke.** Number of genes upregulated (blue), downregulated (red), and unchanged (gray) in mouse astrocytes (mAstro), GABAergic neurons (mGABA), glutamatergic neurons (mGlut), non-neural cells (mNN), and oligodendrocytes (mOligo) in the stroke mouse brain compared to the reference mouse brain.

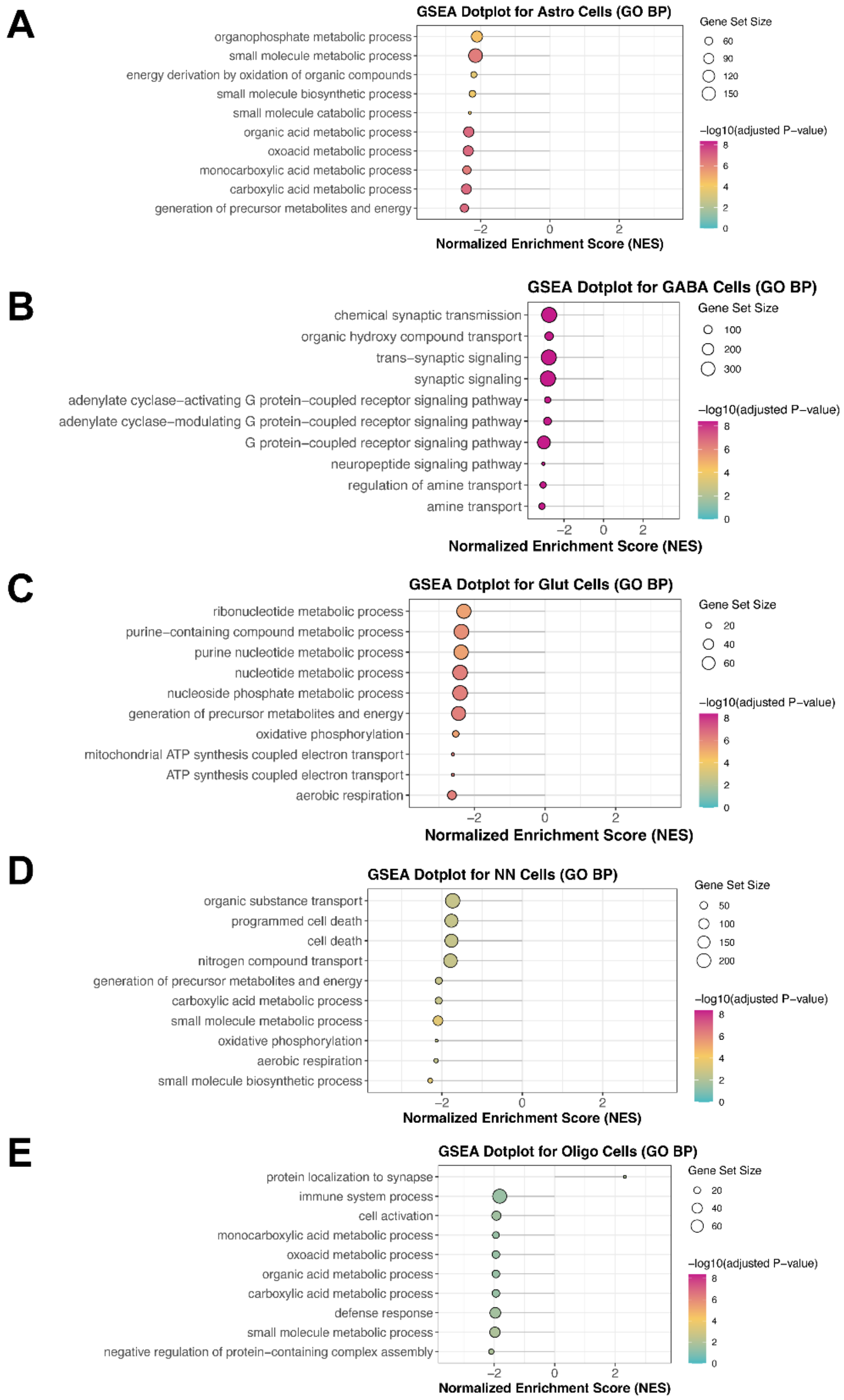


**Suppl. Fig. 19:** Gene Set Enrichment Analysis (GSEA) for different mouse brain cell types in stroke to reference mouse brain dataset. Dot plots show enriched Gene Ontology Biological Processes (GO BP) in (A) astrocytes (mAstro), (B) GABAergic neurons (mGABA), (C) glutamatergic neurons (mGlut), (D) non-neural cells (mNN), and (E) oligodendrocytes (mOligo). The x-axis represents the Normalized Enrichment Score (NES), the color scale indicates the adjusted p-value. Dot size indicates gene set size.

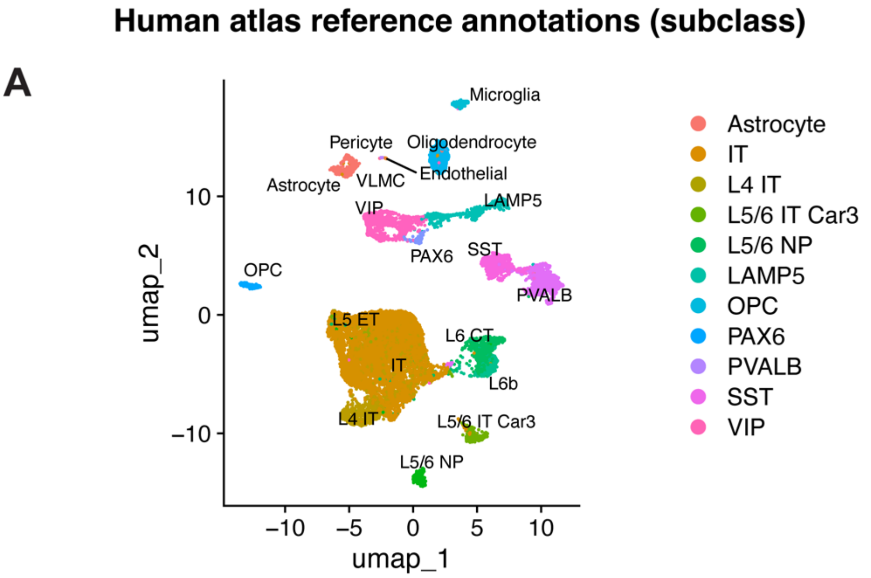


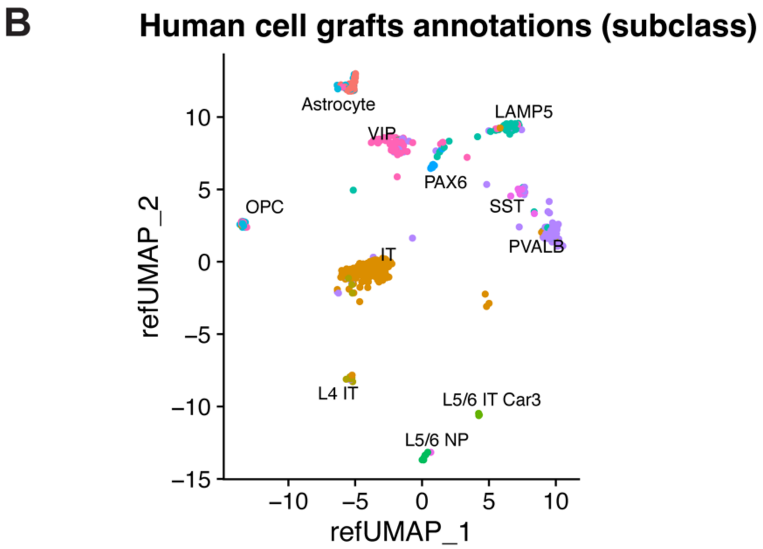


**Suppl. Fig. 20: Identification of cell type subtypes in transplanted cell grafts using human cortex reference atlas**. (A) UMAP of 76,533 nuclei from reference adult human cortex from Allen brain atlas colored by cell type subclass. (B) UMAP of 1149 nuclei from transplanted human cell grafts (n = 5) colored by cell type subclass.

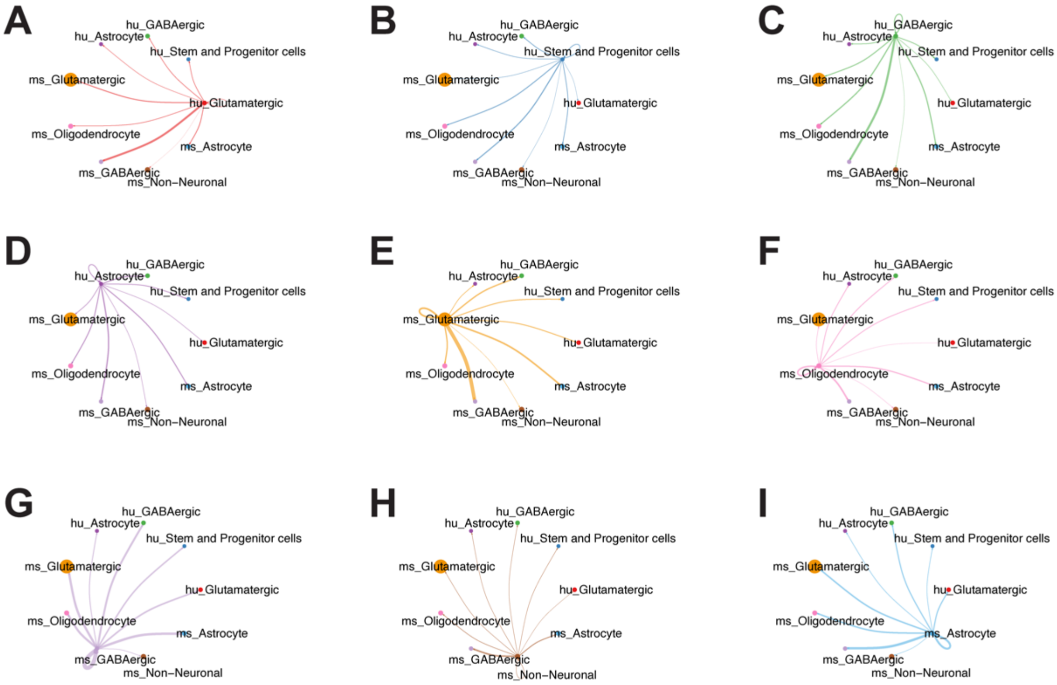


**Suppl. Fig. 21: Total number of cell-cell interactions between individual cell types of graft and host**. Interactions are shown from each cell types (A) human Glutamatergic neurons, (B) human stem and progenitor cells (C) human GABAergic neurons (D) human astrocytes, (E) mouse Glutamatergic neurons, (F) mouse oligodendrocytes, (G) mouse GABAergic neurons, (H) mouse non-neural cells, (I), mouse astrocytes. Circle sizes are proportional to the number of cells in each cell type and line width represents the communication probability.

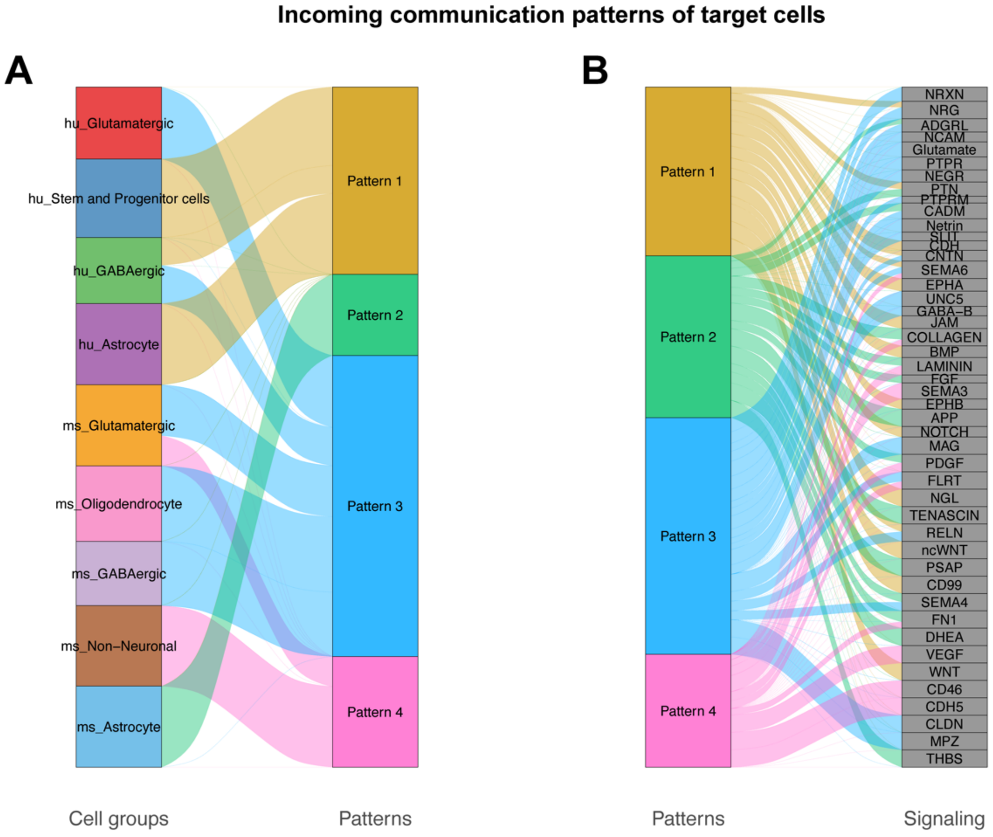


**Suppl. Fig. 22: Molecular communication between cell graft and host**. Riverplot of four incoming interaction patterns for each cell type. The thickness of the flow indicates the contribution of the cell type to each pattern

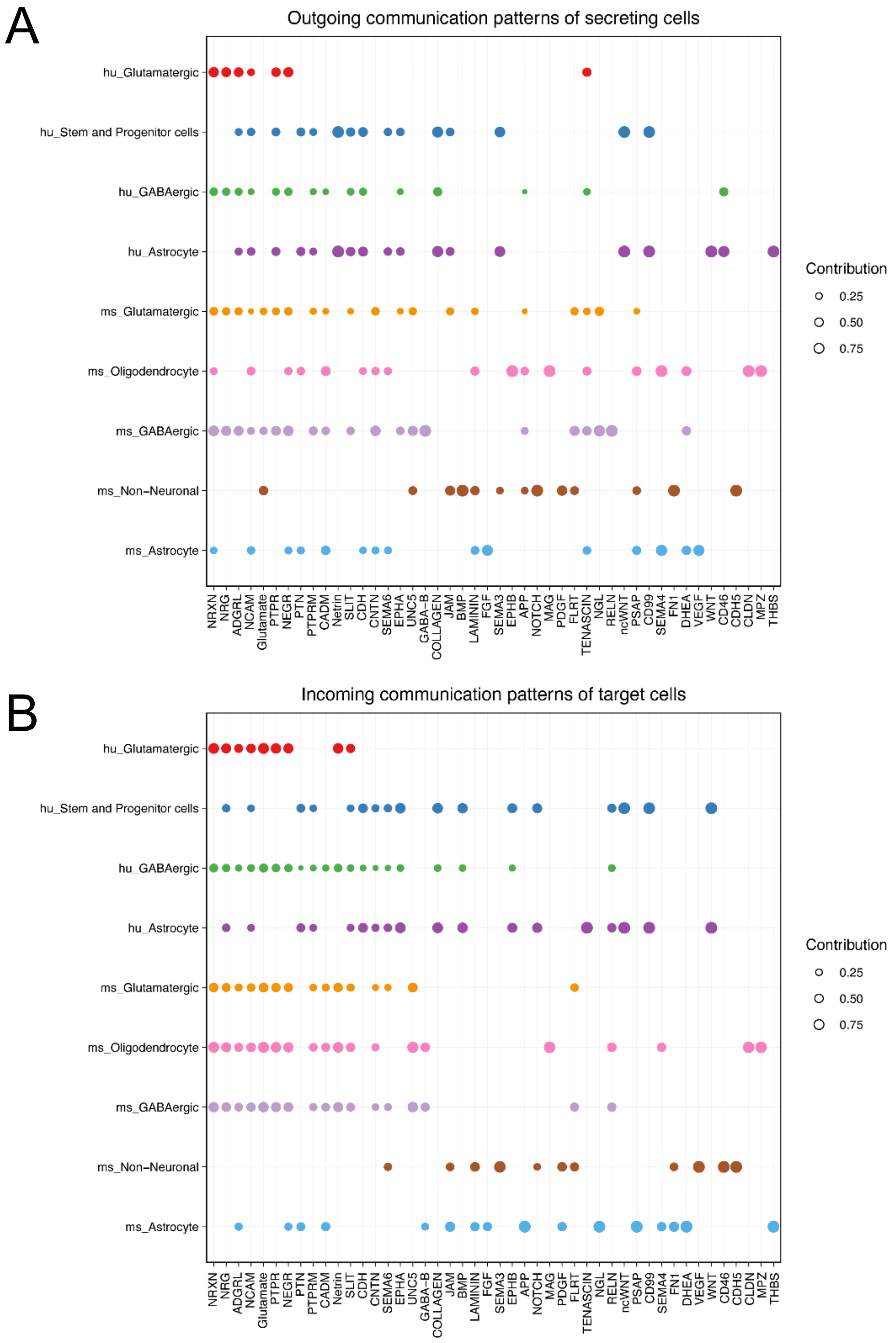


**Suppl. Fig. 23: Signaling pathways in graft-host communication after stroke**. (A) Major pathways that contribute to the sending and (B) receiving signals of individual cell types between graft and host. The dot size represents the calculated communication probability

### interaction of human with mouse cells

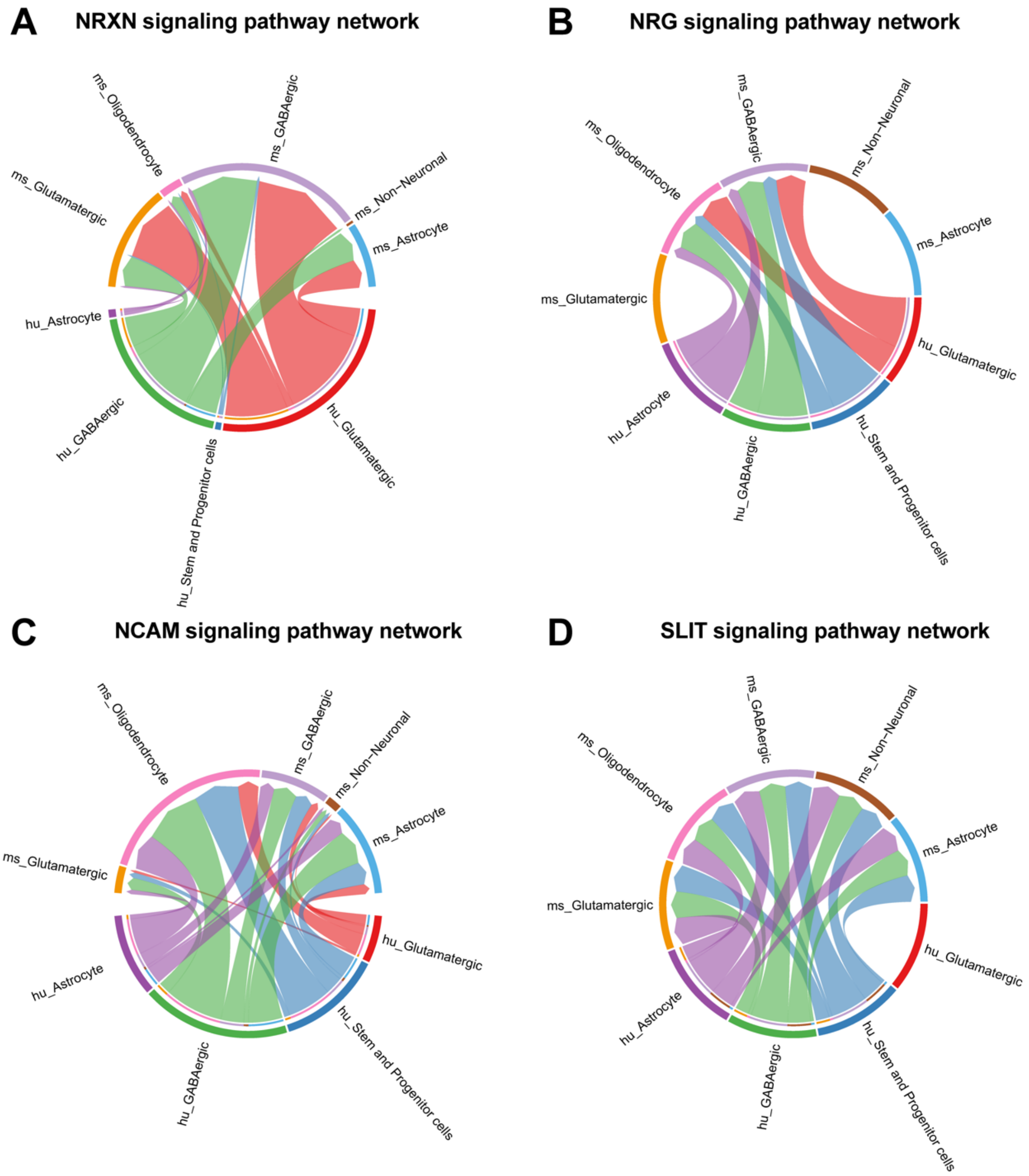


**Suppl. Fig. 24:** Analysis of specific communication pathways between graft and host. Chord plot of neurexin (NRXN), neuregulin (NRG), neural cell adhesion molecule (NCAM), and SLIT showing involvement of individual cell types from graft and host.

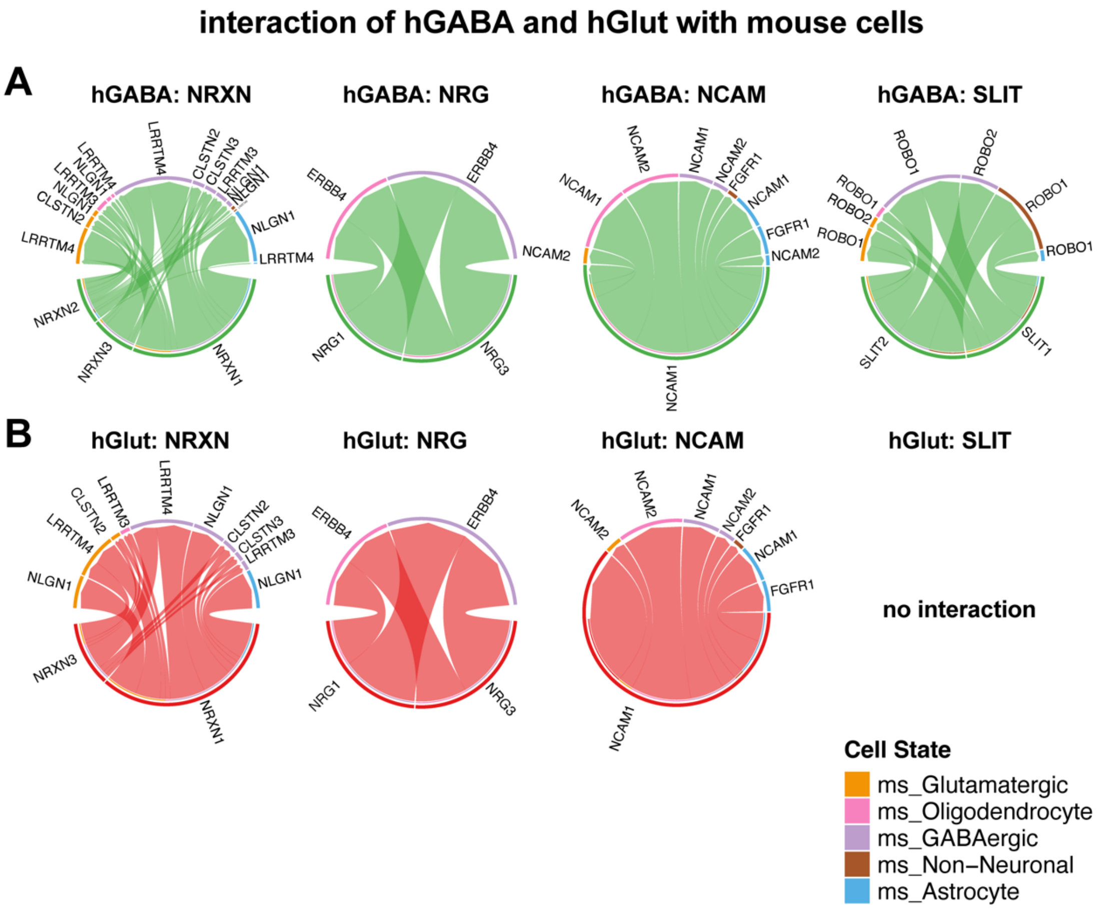


**Suppl. Fig. 25: Analysis of ligand-receptor communication pathways in grafted GABA and Glut neurons with host mouse cells**. (A) Chord plot of neurexin (NRXN), neuregulin (NRG), neural cell adhesion molecule (NCAM), and SLIT signaling showing the most upregulated signaling ligand-receptor for hGABA (green) and (B) hGlut (red).

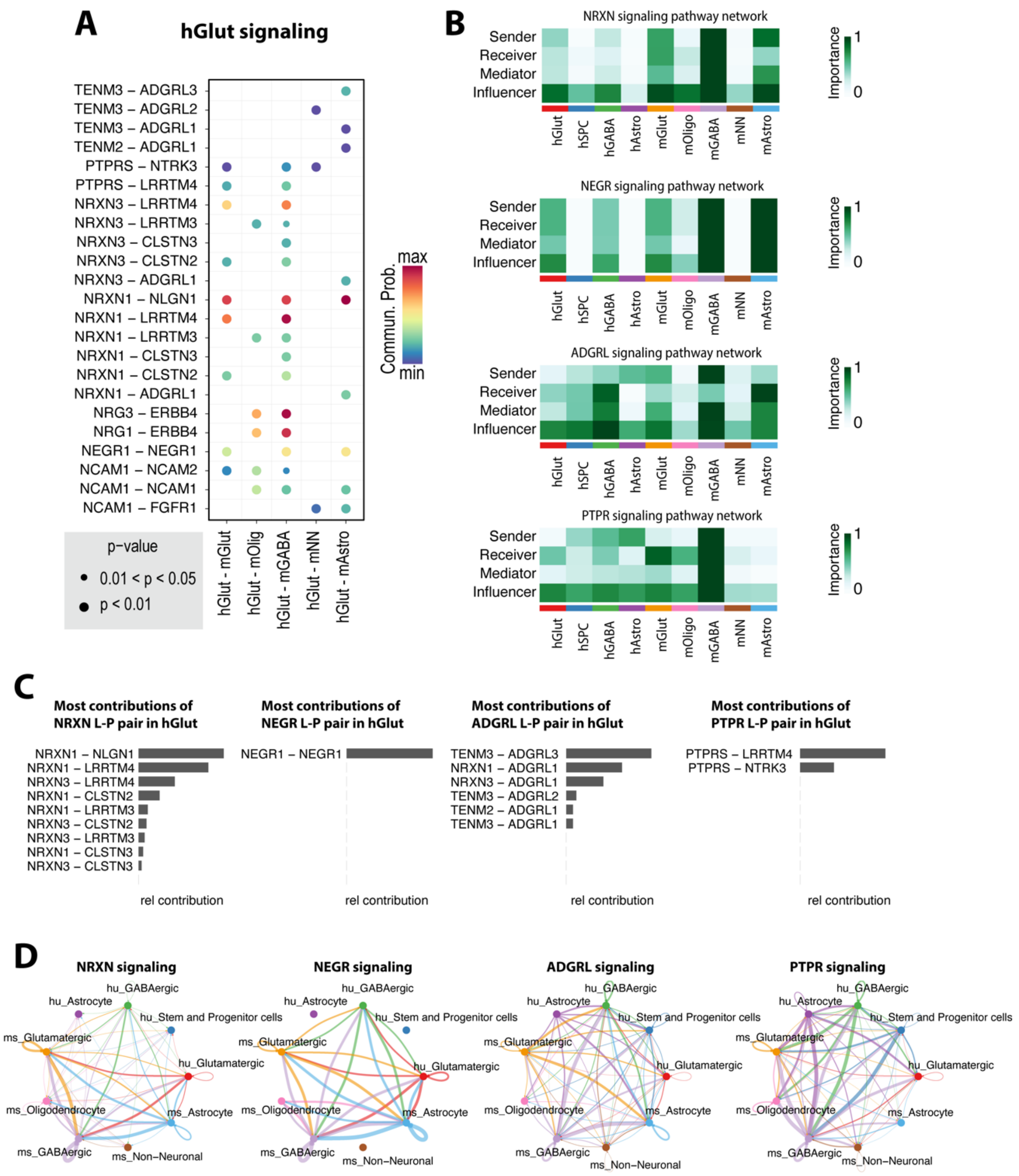


**Suppl. Fig. 26: Molecular crosstalk between Glutamatergic cell grafts and host. (A)** Major ligand-receptor pairs that contribute to the signaling sending from human (hGlut) cells to host cell types from stroked tissue (B) Heatmap show the relative importance of each cell type based on the computed four network centrality measures (sender, receiver, mediator, or influencer) of major hGlut-related pathways: NRXN signaling, NEGR signaling ADGRL signaling, and PTPR signaling. (C) Barplots showing major ligand-partner pairs with most relative contributions in hGlut cells. (D) Number of cell-cell interactions between individual cell types from graft and host for pathways NRXN, NEGR, ADGRL and PTPR.

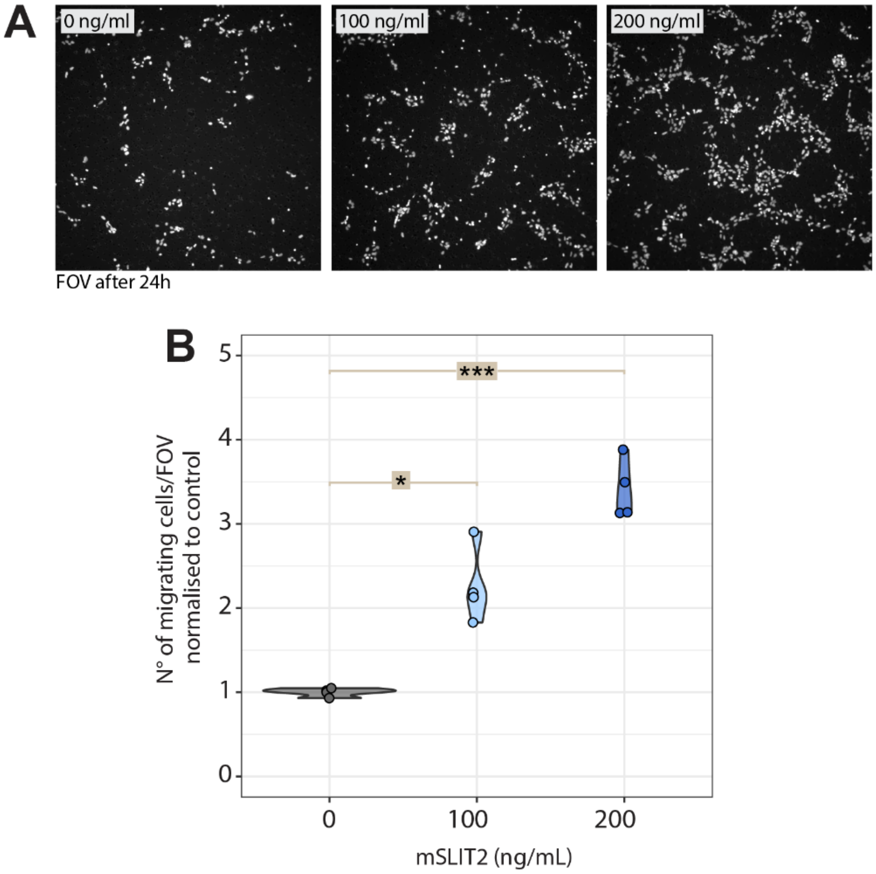


**Suppl. Fig. 27: Migration of hNPCs**. Boyden chamber assays of hNPCs towards 0nM, 100 ng/ml, or 200 ng/ml of recombinant SLIT2 protein. (A) Representative images showing cells that successfully migrated towards different concentrations of SLIT2 protein. (B) Quantification. Violin graphs are plotted as mean ± sem. Significance of mean differences was assessed using an unpaired t-test. Asterisks indicate significance: *$p < 0.05$, **$p < 0.01$, ***$p < 0.001$.
